# Mechanisms of Dominant Electrophysiological Features of Four Subtypes of Layer 1 Interneurons

**DOI:** 10.1101/2022.08.23.505010

**Authors:** John Hongyu Meng, Benjamin Schuman, Bernardo Rudy, Xiao-Jing Wang

## Abstract

Neocortical Layer 1 (L1) consists of the distal dendrites of pyramidal cells and GABAergic interneurons (INs) and receives extensive long-range “top-down” projections, but L1 INs remain poorly understood. In this work, we systematically examined the distinct dominant electrophysiological features for four unique IN subtypes in L1 that were previously identified from mice of either gender: Canopy cells show an irregular firing pattern near rheobase; Neurogliaform cells (NGFCs) are late-spiking, and their firing rate accelerates during current injections; cells with strong expression of the *α*7 nicotinic receptor (*α*7 cells), display onset (rebound) bursting; vasoactive intestinal peptide (VIP) expressing cells exhibit high input resistance, strong adaptation, and irregular firing. Computational modeling revealed that these diverse neurophysiological features could be explained by an extended exponential-integrate-and-fire neuron model with varying contributions of a slowly inactivating *K*^+^ channel (SIK), a T-type *Ca*^2+^ channel, and a spike-triggered *Ca*^2+^-dependent *K*^+^ channel. In particular, we show that irregular firing results from square-wave bursting through a fast-slow analysis. Furthermore, we demonstrate that irregular firing is frequently observed in VIP cells due to the interaction between strong adaptation and a SIK channel. At last, we reveal that the VIP and *α*7 cell models resonant with Alpha/Theta band input through a dynamic gain analysis.

**Significance Statement:** In the neocortex, about 25% of neurons are interneurons. Interestingly, only somas of interneurons reside within Layer 1 (L1) of the neocortex, but not of excitatory pyramidal cells. L1 interneurons are diverse and believed to be important in the cortical-cortex interactions, especially top-down signaling in the cortical hierarchy. However, the electrophysiological features of L1 interneurons are poorly understood. Here, we systematically studied the electrophysiological features within each L1 interneuron subtype. Furthermore, we build computational models for each subtype and study the mechanisms behind these features. These electrophysiological features within each subtype should be incorporated to elucidate how different L1 interneuron subtypes contribute to communication between cortexes.

## Introduction

Neocortical Layer 1, the most superficial layer of the cerebral cortex, is the main target of extensive “top-down” information conveyed by long-range inputs from other cortical regions and subcortical structures (Cauller, 1995; Garcia-Munoz and Arbuthnott, 2015; D’Souza and Burkhalter, 2017; Schuman et al., 2021). The integration of these inputs into local processing is thought to be mediated by a population of GABAergic inhibitory interneurons (INs) residing in L1 in addition to INs in deeper layers with dendrites or axons in L1 (Larkum, 2013; D’Souza and Burkhalter, 2017; Schuman et al., 2021). To date, modeling studies have mostly neglected L1 INs in computational neuroscience. However, in order to study the impact of long-range inputs on cortical functions, a detailed understanding of local L1 circuitry is inevitable (Schuman et al., 2021).

To dissect this process, L1 INs have been classified into different subtypes (Schuman et al., 2019). Not limited to the L1, a classification system considering the definitive features that differentiate IN populations, such as gene expression, electrophysiological properties, and morphology is necessary to study the role of each IN subtype in cortical circuits (Tremblay et al., 2016). Essential findings have been achieved by tagging IN subtypes through genetic strategies (Tremblay et al., 2016), or exploiting the advantages of the Patch-seq technique to glean electrophysiologic and transcriptomic information from the same cell (Gouwens et al., 2020; Scala et al., 2021). In a previous study, we identified four unique populations of INs with somas in L1, each with a distinct molecular profile, morphology, electrophysiology, and connectivity. These four types were Canopy cells, Neurogliaform cells (NGFCs), cells with high levels of the *α*7 nicotinic receptor (*α*7 cells), and vasoactive intestinal peptide (VIP) expressing cells (Schuman et al., 2019).

However, models *in silico* of the functional properties of these cells are still lacking, which hinders the incorporation of these specialized L1 IN subtypes into circuit models. Models of single neurons have proved to be important for understanding the complex behavior and function of neuronal circuits (Traub and Miles, 1991; Llinás, 1988). From recent data (He et al., 2016; Schuman et al., 2019; Goff and Goldberg, 2019; Prönneke et al., 2020), it is of particular interest to understand the irregular firing pattern, which among INs is mostly observed in VIP cells (Tremblay et al., 2016). The irregular firing was a term initially used to describe the unpredictable firing pattern from trial to trial in response to the same depolarizing current injections, a unique feature observed in cells co-expressing VIP and calretinin from the sensory-motor cortex of rats (Cauli et al., 1997), and a subgroup of cannabinoid receptor-1 (CB_1_) positive INs in the somatosensory cortex of mice (Galarreta et al., 2004). Previously, one modeling study suggested the irregular firing arose from the noise-introduced transition between bistable states (Stiefel et al., 2013), which required fast activation kinetics of *K*^+^ channels. In contrast, another study proposed that the clustered spiking depended on the slow inactivation of a Kv1 current (Sciamanna and Wilson, 2011). These hypotheses are worth a re-evaluation in our L1 dataset.

In this study, we identify salient electrophysiological features (e-feature) based on the firing patterns within our L1 IN dataset, namely, irregular spiking (IR), accelerating (Acc), onset bursting (OB), and spike frequency adaptation (Adap). Further, we systematically examine the distribution of these features across the previously identified subtypes: Canopy cells show IR pattern near rheobase; NGFCs are late-spiking, and ACC during current injections; *α*7 cells display OB; and heterogenous VIP cells can exhibit OB, Adap, or IR in the same cell. We then construct generic exponential integrate-and-fire (EIF) models for each IN subtype. We, especially, reproduce IR, OB, and Adap in a generic VIP cell model mimicking a “hero” VIP cell in the dataset. This model includes a SIK channel, a T-type *Ca*^2+^ channel, and a spike-triggered *Ca*^2+^-dependent *K*^+^ channel. A slow-fast analysis shows that irregularity in VIP cells is square-wave bursting. Furthermore, we compare the heterogeneous firing patterns reproduced by varying parameters and conclude that irregular firing is more frequently observed in VIP cells due to the interaction between strong adaptation and the SIK channel. We further test the frequency-dependent response of the VIP cell model through a dynamic gain analysis, and find that the VIP cell model can resonate with Alpha/Theta band input, enabled by the dynamics of the T-type *Ca*^2+^ channel.

## Materials and Methods

### Data

The L1 interneuron data used in this study were published previously in (Schuman et al., 2019), where a detailed description is available. The data were collected from mice of either gender. The four types of interneurons we used in this study are classified based on a combination of gene markers, electrophysical measurements, morphological features, and connectivity. See (Schuman et al., 2019) for details. Within this dataset, the electrophysiological recordings used extremely small increments of positive current injection (1 to 10*pA*) such that the behaviors around the rheobase have great fidelity. For example, the recording of the “hero” VIP cell used 1*pA* as the current step. These fine recordings allow us to study the mechanisms behind different electrophysiological features.

The cells of which the maximum spike count in all the sweeps is less than ten during the 1s current step are excluded in this study. In Figure 1B, Only the cells with a sweep that has 6 APs to 12 APs from 100 ms to 1s are analyzed. If multiple sweeps from the same cell are available, the sweep with the closest AP numbers to 9 is analyzed.

**Figure 1.**
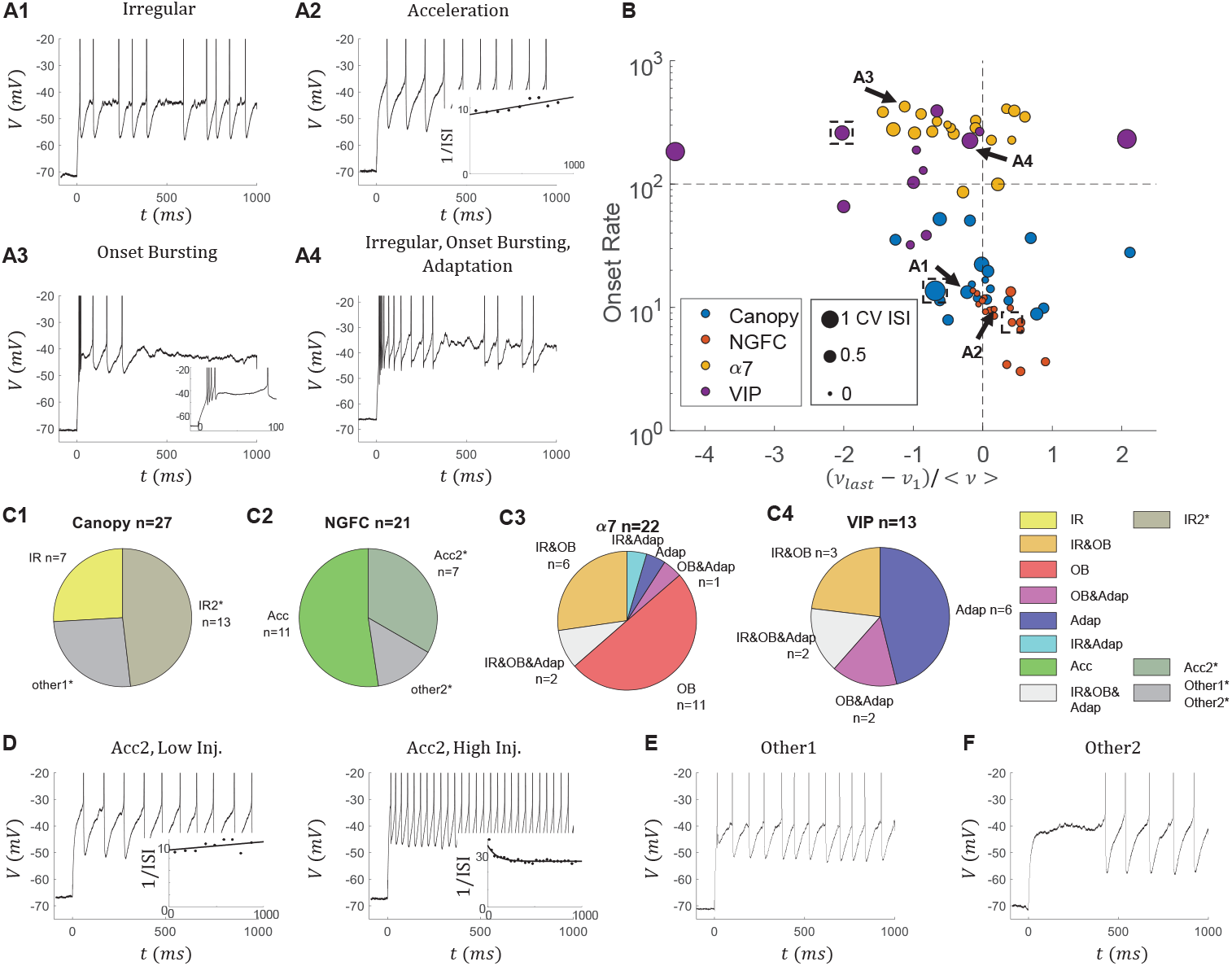
Dominant electrophysiological features of L1 interneuron subtypes. (A) Examples of traces that show four different types of features: irregular firing (IR), Acceleration (Acc) in the firing rate, onset bursting (OB), and adaptation (Adap) in the firing rate. In (A2), the inset shows the 1/ISI and the corresponding fitted line. In (A3), the inset shows the zoomed-in traces during the first 100*ms*. Traces with about 10 APs are selected as examples. (B) Scatter plot of the L1 INs from four subtypes in an E-feature space (*n* = 69). Each dot represents one INs. Only the cells with a sweep that has about 9 APs from 100*ms* to 1*s* are analyzed in this panel (average firing rate < *ν* >= *nAP*/0.9*s* = 10 Hz). The x-axis shows the normalized change of the instantaneous firing rate (*ν* = 1/*ISI*) from the last to the first ISI between 100*ms* and 1*s*. The y-axis is the onset firing rate, calculated by the first ISI. The size of the point represents the CV ISI (> 200*ms*) of that sweep. The arrows show the cells from (A). The dashed boxes indicate the “hero” cells to model except the *α*7 cell. See main texts for details. (C) Distribution of the four features in different subtypes of L1 interneurons. (*) IR2 and Acc2 features are classified by looser criteria than IR and Acc, respectively (See main text). Cells with IR2 or Acc2 are only marked in the Canopy cells and NGFCs for simplicity. 11/27 Canopy cells, 4/21 NGFCs, 9/22 *α*7 cells, and 8/13 VIP cells have an IR2 feature. Notice all VIP cells are classified as having an IR or IR2 feature. 5/27 Canopy cells, 8/21 NGFCs, 0/22 *α*7 cells, and 0/13 VIP cells are classified as having an Acc2 feature. (D) Two sweeps from an exampled cell with an Acc2 feature. The cell shows acceleration in the firing rate when the injection current is low, while adaptation when the injection current is high. (E, F) The cells that do not show any feature are indicated as “other1” or “other2”. The “other2” cells are late-spiking, but the “other1” cells are not.

### AP detection and ISI

The spikes are detected when the voltage trace *V* (*t*) crosses a detection threshold *V_det_*. This threshold is determined as follows: We first set up a upper bound as *V_up_* = min{max(*V* (*t*)), 0*mV* }, then a lower bond as *V_low_* =median(*V* (*t*)). Next, we set the detection threshold as *V_det_* = max{0.8*V_up_* + 0.2*V_low_,* −20*mV* }. The ending detection threshold *V_det_* is within the range −20*mV* to 0*mV*, and it is robust when the AP maximum voltage and AP threshold vary across different cells. The *i*th AP is detected at *t_i_* if *V* (*t_i_*) < *V_thr_* and *V* (*t_i_* + Δ*t*) ≥ *V_thr_*, with the resolution in our recordings Δ*t* = 0.05*ms*. We further exclude any noise-introduced detections within 0.5*ms* following another AP.

We calculate the ISI as ISI(*t_i_*) = *t_i_*_+1_ − *t_i_*. We define instantaneous firing (IF) rate as 1/ISI. In some cases, we fit the curves of instantaneous firing rate to an exponential function *a* + *b* exp(−*cx*) by using the Matlab build-in function *fit*.

### Criteria for electrophysiological features

IR: The IR score and CV ISI are calculated for the sweeps with more than 5 spikes during 500*ms* to 1*s*. The IR score of a pair of ISIs is defined as 1 − min{ISI(*t_i_*)/ISI(*t_i_*_+1_), ISI(*t_i_*_+1_)/ISI(*t_i_*)}, where *t_i_* > 500*ms*. To exclude the potential impact from noise, only the cells with three different ISI pairs, in all the sweeps of the cell, of which the IR score is bigger than the threshold 0.4, are classified as having an IR feature. The IR score of a sweep is defined as the maximum IR score of all ISI pairs of the sweep. CV ISI=*std*(ISI(*t_i_*))/*mean*(ISI(*t_i_*)), where *t_i_* > 500*ms*. If a cell only has one ISI pair, of which IR score is bigger than the threshold during 200*ms* to 1*s*, but is not classified as having an IR feature, the cell is classified as having an IR2 feature. In the model, the IR score and CV ISI are measured from 3*s* to 5*s* during the injection period. The IR width is only measured for the cells with an IR feature. It is defined as the number of sweeps with an IR score bigger than the threshold 0.4 times the current step Δ*I_inj_* of the corresponding recordings.

Acc: We do a linear regression on the curve of 1/ISI for each sweep to calculate the slope. A cell is classified as having an Acc feature if its slopes from all the sweeps are positive. If a cell has more than two positive slopes (at least two sweeps are accelerating) but is not classified as having an Acc feature, the cell is classified as having an Acc2 feature.

OB: We fit the corresponding IF curve to an exponential decay function for the sweeps from a given cell with at least four spikes (three ISIs) during the first 100*ms* of the current injection. If all the fitted IF curves drop at least 50% during this 100*ms*, and if the first ISI in all these sweeps is lesser than 10*ms*, which suggests the onset IF is bigger than 100*Hz* for all these sweeps, the cell is classified as having an OB feature.

Adap: we fit the IF curves of a given cell during 100*ms* to 1*s* to an exponential function *f_adap_*(*t*) for the sweeps with more than six spikes (five ISIs) during this period. The fraction of the dropped firing rate is defined as Adaptation Index (AI= 1 − *f_adap_*(1*s*)/*f_adap_*(100*ms*)). If the fitted IF curve drops at least 20% (AI> 0.2) in 900*ms* for all the sweeps, the cell is classified as having an Adap feature.

### Measurements of electrophysiological properties

The time constant *τ* of a cell is calculated by fitting the voltage trace from the 100*ms* following the end of the sag step to an exponential decay function on each sweep, then averaging across all the sweeps. Resistance *R* is calculated by first dividing the voltage difference (Δ*V*) over the current differences (Δ*I_inj_*) before and during the sag step, then averaging across all the sweeps. Capacity *C* is calculated by dividing the timescale over the resistance in each sweep and then averaging across all the sweeps.

The delay of the first spike is defined as the time of the first spike of the sweep to current injection onset, measured at the sweep with the lowest injection current and at least 2 APs, such that we exclude the possibility that a single spike is triggered by large fluctuation just below the rheobase. We analyze the slope and rheobase of the firing rate curve (f-I curve) by fitting the spike counts during 1*s* to a ReLU function *f* (*I*) = *k* max {*I − I*_0_, 0}, where the resulting *k* is reported as the f-I slope and *I*_0_ as the rheobase. Only the sweeps with lesser than 40*Hz*, or the first sweeps with APs are used to do the ReLU fitting, such that the fitted curve better represents the behavior around the rheobase.

To measure the properties of APs, we exclude the first two APs, which limits the effect of APs during the onset bursting, and combined remained APs from all the sweeps of which the spike count is less than 40. For each AP, the maximum voltage is calculated from the 2*ms* time window following the trace past the AP detection threshold; then, all the APs are aligned by setting *t* = 0 when the voltage reaches the maximum. Maximum rise and decay slopes are calculated during [−1*ms,* 10*ms*]. The AP reset is calculated as the minimum voltage during [0, 10*ms*]. The AP threshold is calculated as the voltage when the voltage deviation is 20*mV/ms*. The halfwidth of AP is calculated as the time of the AP above the midpoint between the maximum voltage and the threshold. Linear interpolation is used in calculating the threshold and halfwidth to improve resolution. Then, all these properties are averaged across all the APs from a cell.

All the measured properties are listed in Table 2. Some values are not identical as in (Schuman et al., 2019) because we exclude cells with fewer than ten spikes.

### EIF models

We implement our models in Python3.9 with the Brian2 package. We use the default ODE solver in Brain2 with a timestep *dt* = 0.1*ms*. Following (Fourcaud-Trocmé et al., 2003), our EIF models have the form

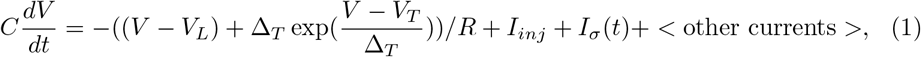

where *V* represents the voltage; *C*, *R* are the passive parameters, capacity, and input resistance; *V_L_* is the reversal potential; *V_T_* is the voltage threshold, and Δ_*T*_ is the curvature at *V_T_*; *σ* is a Gaussian noise term. *I_inj_* is the tonic injection current. *I_σ_*(*t*) = *ση*(*t*) is the fluctuated synaptic or channel current, where *η*(*t*) is an Ornstein-Uhlenbeck process with zero mean and unit variance. For most of the simulation, the correlation time *τ_σ_* of the process is *τ_σ_* = 0.5*ms*, representing an almost-white channel noise. The results are indistinguishable by using a white noise directly. For the simulations in the dynamic gain analysis in Figure 12, the correlation time *τ_σ_* = 5*ms*, representing the fluctuated synaptic noise.

Noticing that the voltage threshold *V_T_*, representing the voltage where the slope is zero, is not the same as the AP threshold measured from the data, where the slope is 20*mV/ms*. All the models have a 2*ms* refractory period after each AP.

In addition, we have included different currents to reproduce the rich dynamics of L1 interneurons.

To reproduce the IR and Acc, we include a SIK current as in (Sciamanna and Wilson, 2011) dynamics:

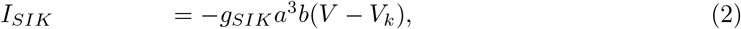

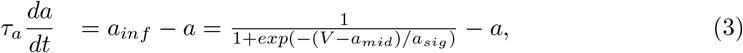

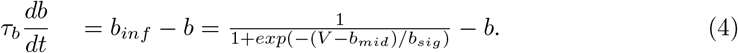

where *V_k_* = −90*mV*, *τ_a_* = 6*ms*, *τ_b_* = 150*ms*, *a_sig_* = 10*mV*, *b_sig_* = −6*mV*, *a_mid_* = −50*mV* + *V_μ_*, *b_sig_* = −65*mV* + *V_μ_*. We shifted the mid point of equilibrium curve *a_inf_, b_inf_* by *V_μ_* = 20*mV* to have the desired window currents. We vary the conductance *g_SIK_* in different models.

To reproduce OB, we include a T-type *Ca*^2+^ current as in (Smith et al., 2000):

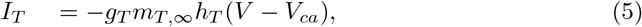

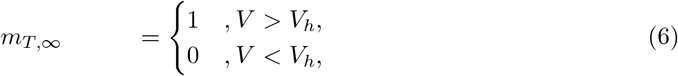

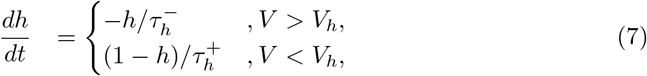

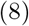

where *V_ca_* = 120*mV*, *V_h_* = 60*mV*, 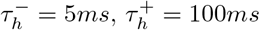.

To reproduce Adap, we include a spike-triggered *Ca*^2+^ dependent *K*^+^ current, as in (Liu and Wang, 2001):

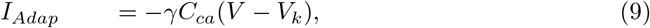

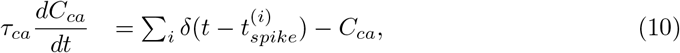

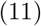

where *C_ca_* represents a dimensionless *Ca*^2+^ concentration in the cell. It jumps by 1 after each spike at 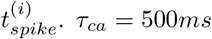.

To tune our model, we first adopt the passive parameters, namely capacity *C* and input resistance *R*, measured from the data. Next, we take the resting voltage as the clamping voltage in the data, which is about −70*mV*. Then, we estimate the reset voltage, threshold, and curvature from the phase diagram of the cell. We also estimate the noise term *σ* based on the standard deviation of voltage below the rheobase. In addition, we consider the change of the gating variable during the APs if a SIK channel is included.

Importantly, we tune the conductance *g_SIK_* of the SIK channel based on the IR score, CV ISI, acceleration, and delay of the 1st spike; we modify the conductance *g_T_* of T-type *Ca*^2+^ based on the firing rate during OB, and we choose the strength of adaptation *γ* based on the AI curve.

We further adjust all the parameters in the model to reproduce all features the best. All parameter values of the model are listed in Table 1.

**Table 1.** Parameters used in the models. The rheobase and f-I slope are measured by fitting the curve of firing-rate at the steady state to a ReLU function.

| | Canopy | NGFC | $\alpha 7$ | VIP |
| --- | --- | --- | --- | --- |
| Resistance $R$ ($M\Omega$) | 86 | 163 | 120 | 400 |
| Capacity $C$ (pF) | 80 | 80 | 80 | 80 |
| Reversal potential $V_L$ (pF) | -67 | -70 | -70 | -69 |
| Threshold $V_T$ (pF) | -38 | -40 | -39 | -40 |
| Reset voltage $V_T$ (pF) | -52 | -52 | -50 | -50 |
| Curvature $\Delta_T$ (mV) | 1 | 1 | 0.5 | 1 |
| Noise Std $\sigma$ (pA) | 20 | 8 | 10 | 10 |
| Conductance $g_{SIK}$ (nS) | 120 | 30 | / | 150 |
| Jump of slow gating variable $\Delta b$ | -0.002 | -0.004 | / | -0.001 |
| Conductance $g_T$ (nS) | / | / | 8 | 10 |
| Adaptation Strength $\gamma$ (nS) | / | / | / | 0.1 |
| Rheobase (pA) | 366.2 | 177.5 | 244.3 | 177.7 |
| f-I slope (Hz/pA) | 1.64 | 0.933 | 1.28 | 0.33 |

**Table 2.**
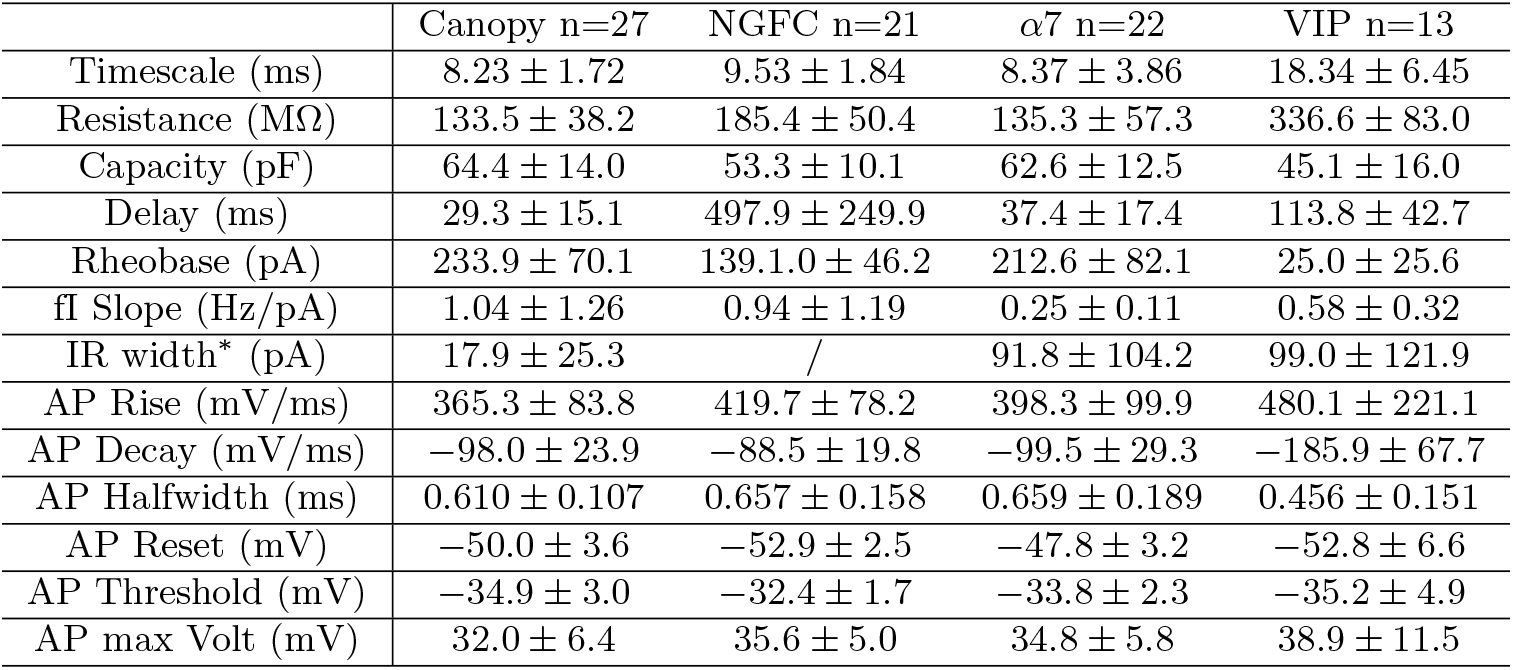
Values are shown as mean ± SEM. Intrinsic electrophysiological properties of the four L1 interneurons subtypes. *: The IR width is only measured for the cells with an IR feature within each subtype (n=7, 9, 5 for Canopy, *α*7, and VIP cells, respectively). See the method for details. The values are not exactly the same as the previous study (Schuman et al., 2019) because we exclude the cells with fewer than 10 APs in the sweep with the maximum injection current.

### Dynamic Gain

The dynamic gain (DG) function *DG*(*f*) is defined as the ratio of the response and the input at a specific frequency. We followed the exact method as described in (Ilin et al., 2013). Here, the response firing rate function is defined as a summation of delta functions at spike time *t_i_*:

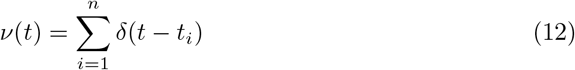

As proposed by (Higgs and Spain, 2009), we calculated the dynamic gain function in the Fourier space as

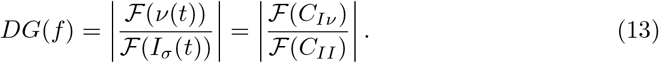

The numerator is the Fourier transformation of input-output correlation. Since the firing rate *ν*(*t*) is a sum of delta functions, *C_Iν_* can be simplified into the averaging current in the time windows around each spike:

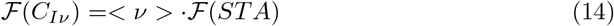

where *STA* represents the spike-triggered average current. We use a 1*s* window and frequency-dependent Gaussian filters to suppress noise.

Since the *I_σ_*(*t*) is a standard OU process, the denominator can be calculated theoretically (see also (Zhang et al., 2022)), as

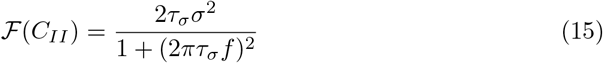

At last, the DG is normalized by the value at 1 Hz *DG*(1).

The 5% confidence threshold is calculated by bootstrapping as in (Borda Bossana et al., 2020). We first generated 500 surrogated response *ν_i_*(*t*) trials by shuffling the original ISIs. Then we calculated the dynamic gain functions *DG_i_*(*f*) of these surrogated responses. The confidence curve is calculated based on each frequency’s mean and variation from the *DG_i_*(*f*).

To calculate the high-frequency profile, we follow (Borda Bossana et al., 2020). The cut-off *f*_0_ is defined as the DG dropps to 70% *DG*(1). To describe the exponential decay of DG at large frequencies, we fit the *DG*(*f*) between *DG*(*f*_0_) and 0.2*DF* (*f*_0_) to *y* = *F* (*f*) = *bf ^−α^*, where *α* represents the decay constant.

### Experimental design and statistical analysis

We analyzed the original pClamp data in Matlab (MathWorks) using a self-developed package. The developed toolbox during this project will be fully available online upon acceptance. The models in this study were developed in Python 3.9.7 (Python Software Foundation) with Brian2 package (Goodman and Brette, 2008), and will be uploaded to the ModelDB upon acceptance. All statistical analysis was performed in Matlab.

The full dataset is available upon request.

## Results

### Identifying dominant electrophysiological features of L1 interneuron subtypes

Previously, we separated the L1 IN population into Canopy cells, NGFCs, *α*7 cells, and VIP cells based on a combination of genetic markers, morphology, connectivity, and electrophysiological features such as input resistance, sag, and latency to 1st spike (Schuman et al., 2019).

Here, we systematically studied electrophysiological features of firing patterns based on the analysis of all the sweeps in our dataset. We first identified four features of interest in this L1 dataset (Figure 1A). Following the Petilla terminology (Ascoli et al., 2008), these features are irregular firing behavior (IR), acceleration of the firing rate (Acc), onset bursting (OB), and adaptation (Adap) of the firing rate. To show how salient these features exist in our dataset, we plot each cell in an e-feature space (Figure 1B). To compare across cells, we used the sweep where the firing rate is about 10*Hz* during the 100*ms* to 1*s* of the current step. The IR is represented by the CV ISI (size), the Acc and Adap features are represented by the normalized change of the firing rate (x-axis), and the OB is characterized by the onset firing rate (y-axis). Though this is a useful tool, it is limited in two ways. First, when the cell fires irregularly, the change in the firing rate may reflect the irregularity but not the trend in the firing rate. Second, the information that resides outside the 10*Hz* sweeps is lost in this analysis. Still, different IN subtypes (colors) are separated in this space. Canopy cells show IR in many cells but no other features; NGFCs only show Acc; *α*7 cells are OB without exceptions; and VIP cells are heterogeneous but mostly show IR and Adap. The traces in Figure 1A are marked by arrows in Figure 1B.

Next, we set quantitative criteria based on all the sweeps, but not just one sweep, to better determine whether each cell shows these four features (Figure 1C). Here, we include more cells than in Figure 1B, since having a sweep of 10*Hz* is not required. Traditionally, irregularity is used to describe a unique, unpredictable firing pattern (Galarreta et al., 2004). This can be further reflected in the abrupt change of the inter-spike intervals (ISI) (Figure 1A1). To quantify it, we define an IR score of a pair of neighboring ISIs as one minus the ratio of the smaller ISI and the larger ISI. If one cell has at least three pairs of ISIs during the last 500*ms* of the depolarizing pulse that have an IR score bigger than the threshold of 0.4, we consider the cell as having an IR feature. We find the IR score is more robust than the traditional measurement CV ISI in quantifying the irregular firing for the following reasons. On the one hand, we have fine recordings around the rheobase, such that we can collect ISI pairs from different sweeps to distinguish the irregular firing from pure noise. On the other hand, many L1 INs show Acc or Adap, which is hard to distinguish from IR by CV ISI.

We only use the last 500*ms* to avoid the potential false-positive cases because cells with strong adap or OB can change their firing rate rapidly at the onset of the current injection period. We also define a looser criterion for having an IR2 feature, i.e., if a cell does not display an IR feature but has at least one pair of ISIs during the latter 800*ms*. If a cell with an IR2 feature may have an IR feature if the recordings were done with finer current steps, but it may not if the ISI pairs with large IR scores are purely introduced by noise. IR is widely observed in L1 INs except for NGFCs (Figure 1B, C). Noticeably, all the VIP cells have an IR or IR2 feature.

We consider a cell as having an Acc feature if it shows acceleration in the firing rate in all the sweeps. A looser criterion, which only requires one cell to have at least two sweeps around rheobase to show the acceleration in the firing rate, is applied to consider a cell as having an Acc2 feature. These cells with an Acc2 feature usually show acceleration when the injection current is low, but adaptation when the injection current is high (Figure 1D). Cells with an Acc feature are uniquely observed in NGFCs. Further, most NGFCs are considered as having an Acc or Acc2 feature (Figure 1C2), and some canopy cells are found to have an Acc2 feature (5/27).

Then, to distinguish OB and Adap, we define OB and Adap based on different time frames during the current step. We consider a cell to have an OB feature if, first, the onset firing rate is larger than 100*Hz* (first ISI is lesser than 10*ms*) in all the sweeps with at least five APs; and second, the firing rate drops at least 50% in the first 100*ms* in all the sweeps. We consider a cell to have an Adap feature if the firing rate drops at least 20% from 100*ms* to 1*s* in all sweeps. The OB feature is observed in most of the *α*7 cells (20/22, the two *α*7 cells without OB features are just below the OB cut-off in Figure 1B), and some of the VIP cells, while Adap is observed in most of the VIP cells and some of the *α*7 cells (5/22, Figure 1C3, 4).

The cells with no associated features are marked with “other1” or “other2”. While the “other1” canopy cells show no signature (Figure 1D), the “other2” NGFCs are all late-spiking (Figure 1E). The passive parameters are listed in table 2 and shown in Figure 2. The detailed characterizations of individual cells are listed in Extended Data Table 1-1.

**Figure 2.**
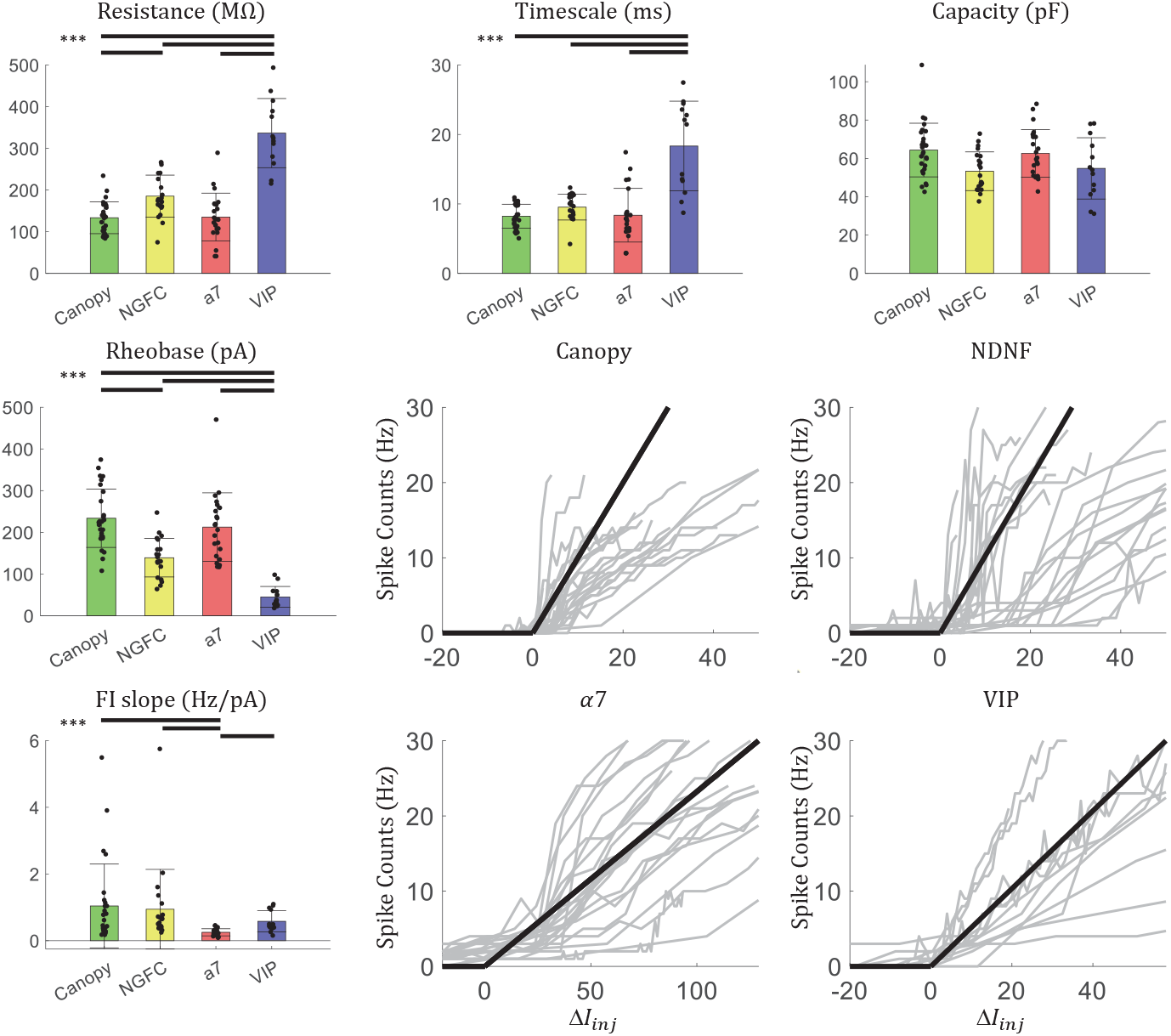
Electrophysiological features of the L1 interneurons. The bars above suggest the corresponding pair is significantly different (*p* < 0.001, Mann-Whitney U test). Rheobase and FI slope are measured by fitting the spike-count curve to a ReLU function. The bold black lines show the ReLU function with the mean FI slope. Grey lines represent individual spike-count curves of individual cells.

In this study, we want to have parsimonious models that reproduce the spectrum of these behaviors and help understand the mechanisms behind these behaviors. To achieve these, instead of tuning our model based on the averaged features of a subtype, we choose one “hero” cell from each subtype that only shows one type of the electrophysiological feature of interest and reproduce the behavior of that cell. However, readers should be aware that our model does not consider heterogeneity within each cell type. The analysis on individual cells is provided in Extended Data Table 1-1 if the heterogeneity itself is of interest. Canopy, NGFC, and VIP “hero” cells are marked by the dashed boxes in Figure 1B. The closest sweep of *α*7 “hero” cell has 17 APs after 100*ms*, with CV ISI= 0.09, *dν/ < ν* >= 0.18, and onset firing rate = 235.3*Hz*.

We do this because the averaged “cell” may not be physically plausible due to the large heterogeneity within an interneuron subtype. We choose to make individual neuron models based on the exponential integrate-and-fire (EIF) neuron (Fourcaud-Trocmé et al., 2003), but not Hodgkin-Huxley models, to minimize the number of parameters we need to tune. In the following sections, we, first, reproduce IR in a canopy cell model; second, capture the Acc and delayed spiking in a NGFC model; third, generate the OB in an *α*7 cell model. Finally, we model a VIP cell that shows IR, OB, and Adap simultaneously.

### Irregular firing is reproduced in a Canopy cell model with a SIK channel

Since canopy cells with an IR feature do not have any other of the features under consideration, modeling IR on canopy cells does not need to consider the potential interactions between different features (Figure 1C1). From the selected “hero” canopy cell. We observed substantial subthreshold fluctuations around −40*mV* from the voltage traces just above the rheobase (Figure 3A, C). This is more obvious in the phase diagram of the cell (Figure 3B, D), where we plot the deviation of the voltage, which is proportional to the net current into the cell, over the voltage. A quasi-stable point is observed around −40*mV* where the net current goes below 0, suggesting some outward current is activated during this range.

**Figure 3.**
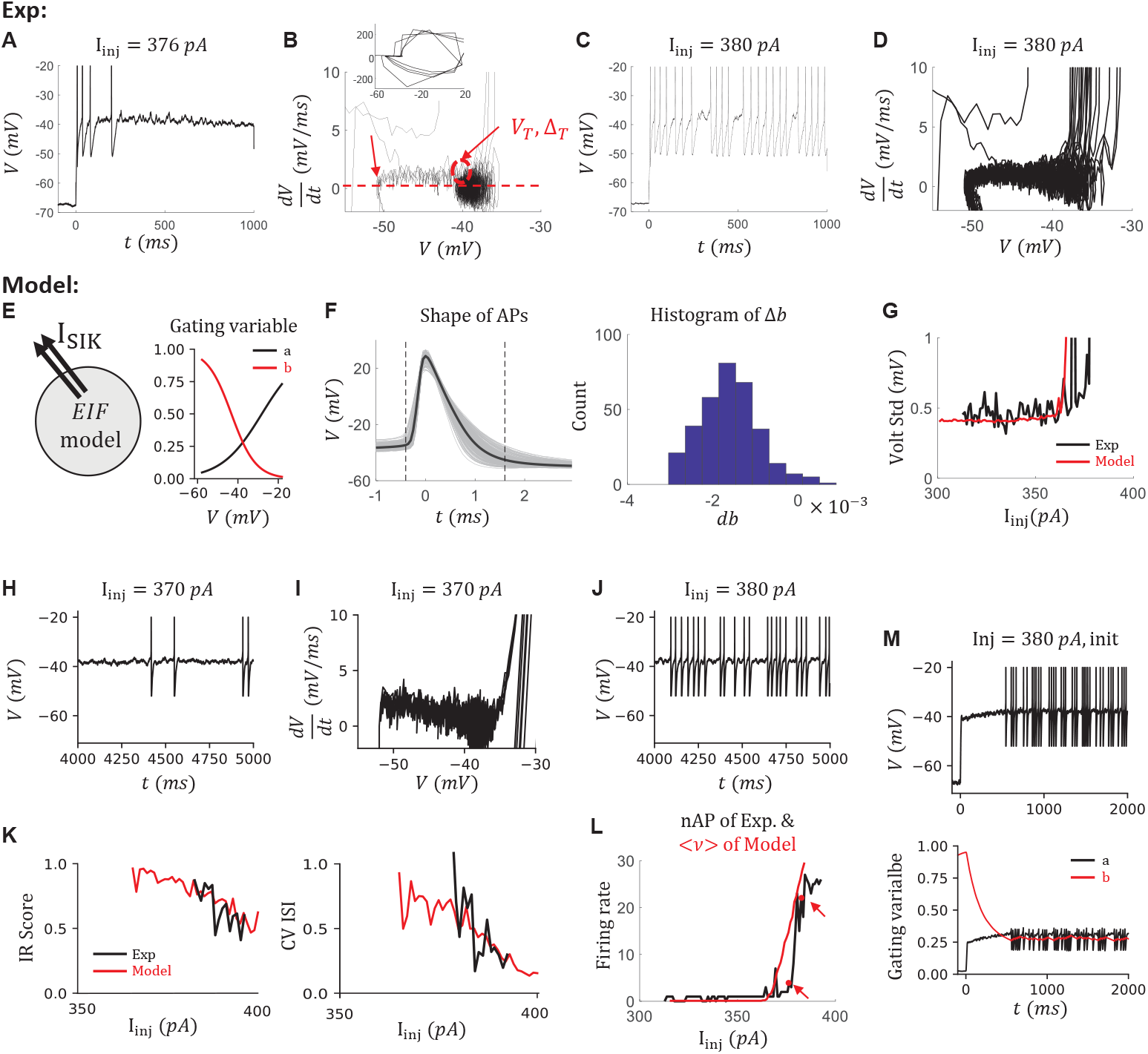
Modeling irregular spiking of a canopy cell with a SIK channel. (A) the voltage trace of an example canopy cell at *I_inj_* = 376*pA*. (B) Phase diagram of the cell. The inset shows the corresponding zoom-out diagram. The y-axis is the slope of the voltage trace, which is proportional to the net current to the cell. Red arrows indicate where we estimate the value of reset, curvature, and firing threshold of the corresponding model. (C, D) the trace and the phase diagram at *I_inj_* = 380*pA*. (E) The sketch of the canopy cell model (left) and the equilibrium value of the gating variables of the SIK channel (right). (F) Δ*b* estimation. Left: the shapes of all the action potentials (AP) from all the sweeps of the same cell. We exclude the first two APs in every sweep. APs are aligned at the maximum of the voltage. The dashed lines indicate *t* = −0.4, 1.6*ms*, respectively. Voltage traces of these 2*ms* time windows are used to calculate the jump of the slow inactivation variable Δ*b*. Right: the histogram of Δ*b* when assuming *b_init_* = 0.28 just before the AP. The value *b_init_* = 0.28 is the average value after 3*s* of injection in the simulation. The average change is < Δ*b* >= −0.0020. (G) The voltage standard deviation measured between 500*ms* to 1*s* during the current injection from the data (black) and the model (red). (H, I) The voltage trace and the phase diagram of the Canopy cell model at *I_inj_* = 370*pA* (J) The voltage trace at *I_inj_* = 380*pA*. (K) The comparison of irregularity from the data and the model by IR score (left) and by CV ISI (right). (L) The comparison of firing rate from the data, estimated by the number of APs during the 1*s* current injection, and from the model, estimated by the steady-state firing rate. The red arrows, *I_inj_* = 376, 380*pA*. (M) The initial dynamics are not considered in this model. Top: the voltage trace that includes the initial transient part when *I_inj_* = 380*pA*. Bottom: the transient dynamics of gating variables of the SIK channel. The black and red lines indicate the fast and slow variables *a* and *b*, respectively.

To mimic the IR at the latter 500*ms* during the current injection, we include a slow inactivating *K*^+^ (SIK) channel in an EIF model (Figure 3E, left). The major parameters of an EIF model, namely reset voltage, firing threshold, and curvature at the firing threshold can be estimated from the phase diagram directly (Figure 3B, red arrows). The slow-inactivation gating variable from the SIK channel can introduce the clustered spiking, which has been known (Wang, 1993). Here, the model of the SIK channel follows the model of a Kv1 channel in PV+ interneurons (Sciamanna and Wilson, 2011), which contains one fast activating gating variable *a* and one slow inactivating gating variable *b*. We shift the equilibrium value of the gating variables to a more depolarized value such that the window current of this channel is around −40 mV (Figure 3E, right). As argued in (Sciamanna and Wilson, 2011) Fig. 8, this is plausible by taking into account that the Kv1 channels are localized on the axon initial segment but not on the soma.

To tune the model, we further consider the change of the gating variable during the APs. We can estimate the change of the slow variable Δ*b* after each AP by solving the differential equation of *b* while replacing the voltage term by the shape of APs (Figure 3F). The initial value of *b* is chosen based on our simulation. This is plausible because the dynamics of *b* are much slower (time constant *τ_b_* = 150*ms*) than the time window of APs (refractory period of the model 2*ms*). Choosing different Δ*b* in the model has little impact on the simulation results. Since the jump of the fast variable Δ*a* doesn’t change the dynamics of the model due to the rapid convergence to the equilibrium point (*τ_a_* = 6*ms*, Figure 3M), we set Δ*a* ≡ 0. Also, we estimate the noise term *σ* based on the standard deviation of voltage before or around the rheobase current during 500*ms* to 1*s* of the depolarization current step (Figure 3G). Importantly, we tune the conductance *g_SIK_* of the SIK channel based on the IR score curve. At last, we adjust all the parameters in the model to best reproduce other cell features, namely the rheobase, the slope of the f-I curve, and the firing behavior of individual sweeps.

The resulting model can reproduce the IR behavior observed in the cell (Figure 3H to J) and the changing of IR behavior when the injection current varies (Figure 3K). The firing rate is also comparable between the data and the model (Figure 3L).

However, since we focus on the irregularity in this section, the dynamics around the beginning of the current injection are not modeled (Figure 3M). As a result, the model shows a delayed-firing feature (Figure 3M, top), because of the slow inactivation of SIK channel (Figure 3M, bottom).

### Mechanism of irregular firing in the Canopy cell model

In our model, The IR feature is reproduced by including a SIK channel. To understand the mechanism, we did a fast-slow analysis on our model (Figure 4). Excluding the noise term by setting *σ* = 0, we observe clustered spiking in the model (Figure 4A). We further assume the slow gating variable *b* is a constant and analyze the dynamics of the fast manifold defined by the voltage *v* and the fast activation variable *a*. As expected for the EIF model, the system has one stable state (SS) and one unstable steady state (US) when the injection current is low or the outward current is large (Figure 4B). The SS branch and US branch emerge through a saddle-node bifurcation on an invariance cycle (SNIC) by increasing *b* over the bifurcation point *b_min_*, (Figure 4C). In the simulation of the full system (Figure 4A, C), the slow variable *b* changes across the bifurcation point *b_min_* over time. Thus the full system oscillates between a resting state (the global SS exists) and a spiking state (the global SS vanishes). This dynamic is known as square-wave bursting (Rinzel and Ermentrout, 1998; Wang and Rinzel, 1995; Rinzel, 1987). Adding noise to the system *σ* > 0, the oscillation between the resting and spiking states mimics the IR firing patterns.

**Figure 4.**
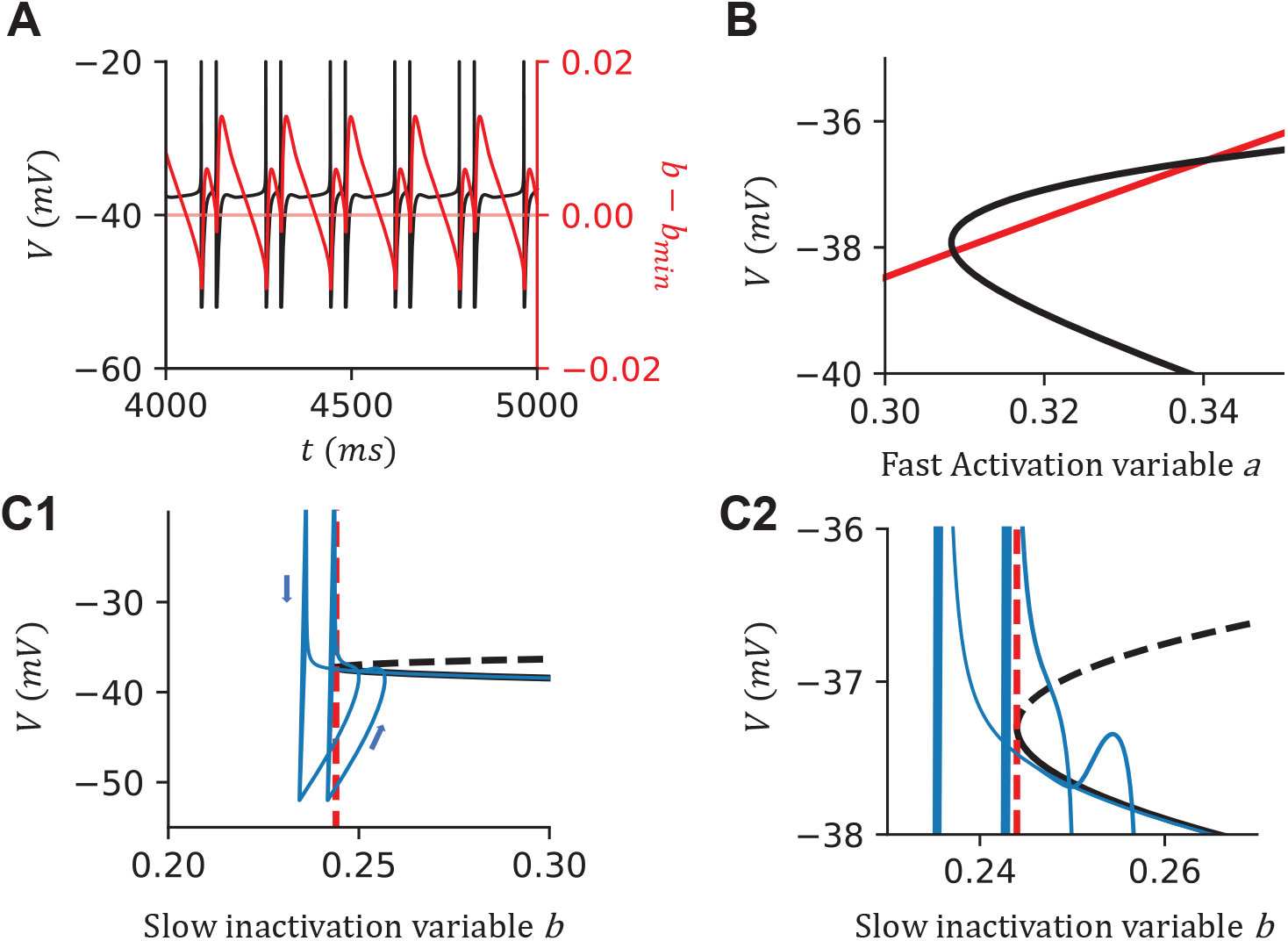
A fast-slow analysis of the Canopy cell model shows that the irregularity results from the square-wave bursting. (A) Sample simulation when *I_inj_* = 375*pA* without noise. The black line indicates the voltage trace, and the red line indicates the difference between *b* and *b_min_*. The red dashed line is at *b − b_min_* = 0, suggesting the boundary between the resting state and the firing state of the cell. (B) Nullclines when *I_inj_* = 375*pA*, *b* = 0.27. The intersections indicate the fixpoints of the fast manifold. (C) The bifurcation diagram when varying *b* with *I_inj_* = 375*pA*. The system undergoes a saddle-node bifurcation on an invariant cycle (SNIC). The black solid and dashed lines indicate the stable branch and the unstable branch, respectively. The blue line indicates the simulation of the canopy cell model while we remove the noise term *σ* = 0. The red dashed line indicates the minimum *b* that the fast manifold has a global SS. The arrows indicate the direction of the dynamics. (C2) shows the zoomed-in dynamics around the bifurcation point.

### Spike frequency acceleration and delayed firing are reproduced in a NGFC model

After capturing the dynamics of IR, we next model the acceleration observed in a NGFC. As shown in Figure 5, the Acc is accompanied by the late onset of spiking (Figure 5B, C). Similarly as in tuning a Canopy cell model, we further characterize the cell based on the corresponding phase diagram (Figure 5D) where a similar quasi-stable point is observed around −40*mV*, the jump of change of the slow variable Δ*b* after each AP (Figure 5E), and the noise level (Figure 5F). Importantly, the firing rate slowly accelerates over the 1*s* injection period (Figure 5G, H).

**Figure 5.**
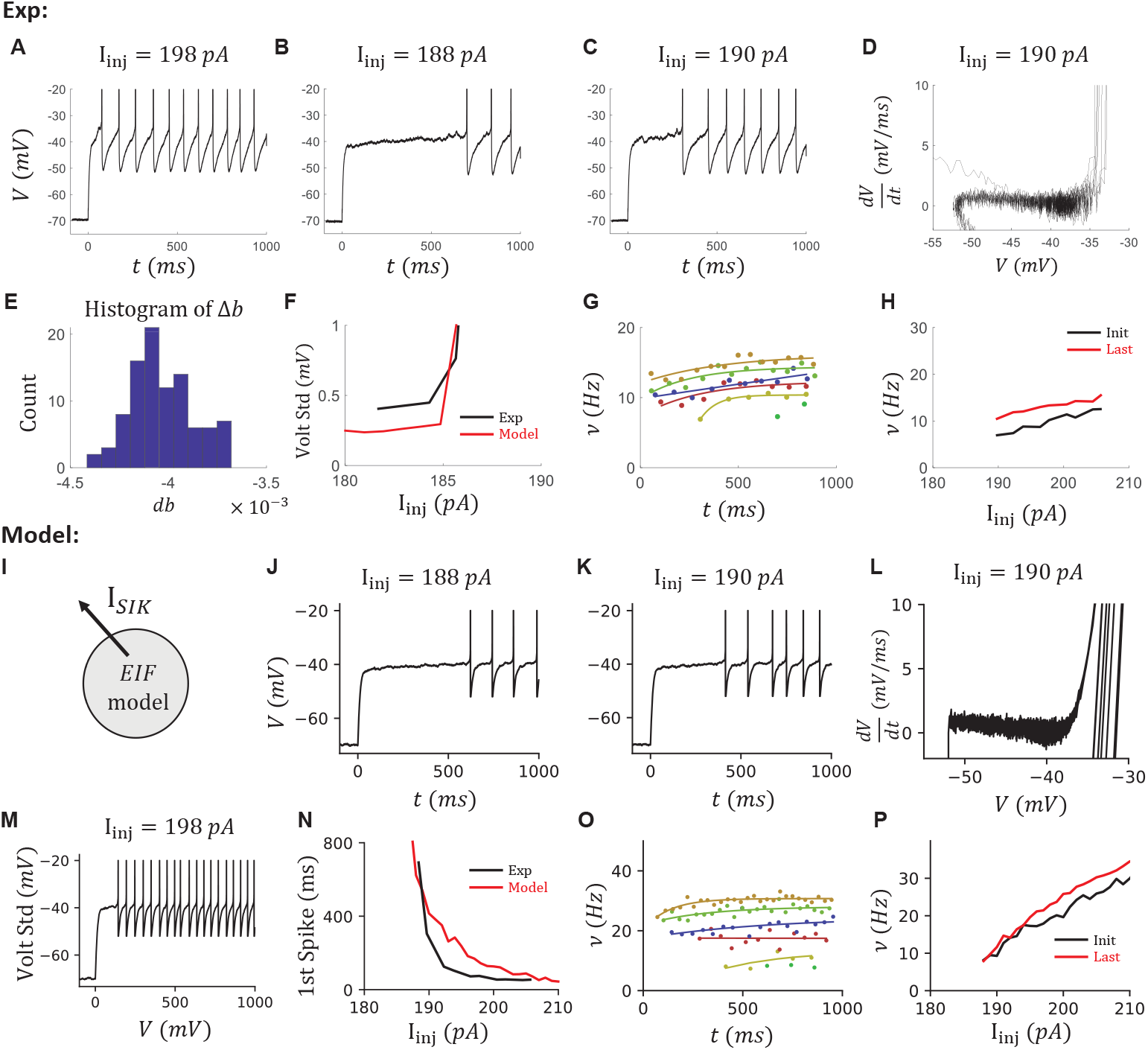
Modeling the acceleration and late-spiking of a NGFC with a smaller SIK conductance. (A to C) Voltage traces of the selected NGFC at *I_inj_* = 198, 188, 190*pA*. (D) The phase diagram of the cell at *I_inj_* = 190*pA*. (E) The histogram of Δ*b* if assume *b_init_* = 0.4 before the AP. The average change is < Δ*b* >= −0.0040. (F) The voltage standard deviations of the data and the model. (G) The instantaneous firing rate *ν* = 1/ISI over time. We fit the ISIs (dot) to an exponential decay curve (line) for different injection currents *I_inj_* = 188, 190, 194, 198, 202, 206*pA* (color). (H) The initial firing rate (black) and the last (red) firing rate at different injection currents. (I) The sketch of the NGFC model. (J to M) Model results corresponding to (B to D, A). (N) The time of the first spike decreases fast when increasing the injection current in both the data and the model. (O, P) The instantaneous firing rate *ν* increases over time in the model. Organized as (G, H)

Since the delayed firing is associated with the SIK current (Storm, 1988), along with our observations, we tune an EIF model with the same SIK channel that has the identical dynamics as in the Canopy cell model to reproduce the ACC feature (Figure 5I). Compared to the Canopy cell model, we reduce the SIK conductance *g_SIK_* to avoid the IR. The relationship between SIK conductance *g_SIK_* and the IR will be discussed later in Figure 11. After a similar tuning procedure, the resulting model is capable of reproducing the Acc feature observed in a NGFC (Figure 5J to P). In addition, the time of the first AP drops fast when increases the injection current, and our model is capable of reproducing this behavior (Figure 5N).

The delayed firing and the acceleration come from the slow inactivation gating variable *b* of the SIK current. To directly show that, we simulate the model with two sequential voltage steps and current steps with a variable resting time Δtime (Figure 6A, B). During the first voltage step (Figure 6A), the model neuron is clamped at −40*mV*. The fast-activating gating variable *a* opens first and increases the SIK current. Then, the slow-inactivating variable *b* gradually closes and decreases the SIK current. The difference between the onset and steady SIK current is about 15*pA* in our NGFC model. During a rheobase current step (Figure 6B), the model neuron depolarizes to a subthreshold voltage but is held there by the fast-activating SIK current. It is only released to fire when the SIK current inactivates enough. During a current step that is above the rheobase, the SIK current also inactivates gradually, which leads to an acceleration of the firing rate. Further, the strength of the SIK current is subjected to the initial state of the slow variable *b*. If the second voltage step or current step is close to the first, the slow variable *b* does not fully restore to the open state. As a result, the peak of the SIK current and the delay of the first spike is smaller in the 2nd step compared to the 1st step (Figure 6C).

**Figure 6.**
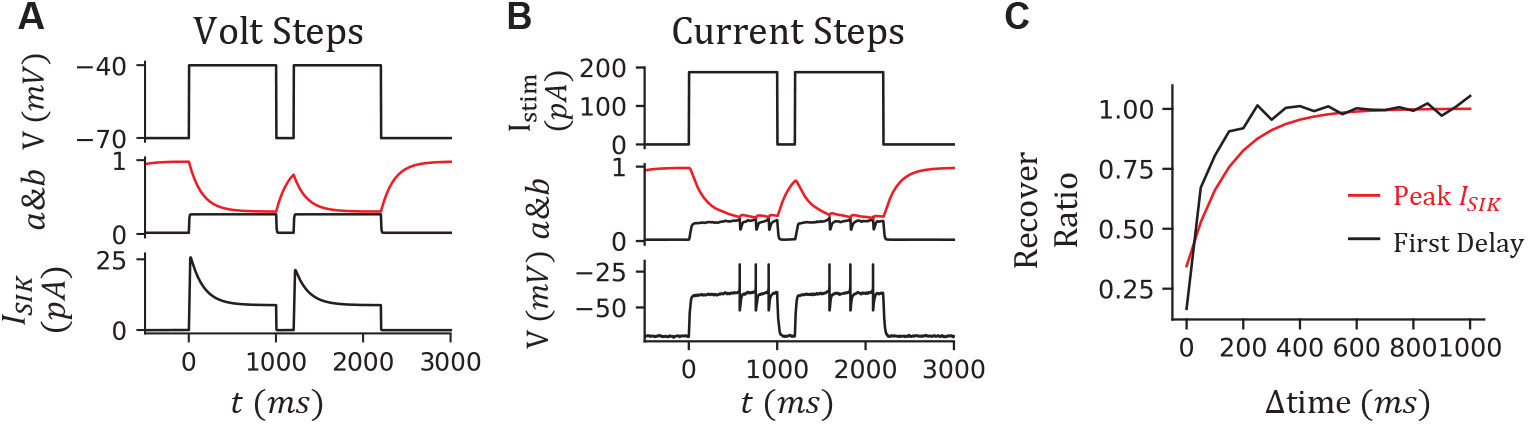
Recovering dynamics of the NGFC model. (A) Simulation of the two voltage steps *V_clamp_* = −40*mV* with a resting time Δtime = 200*ms*. From top to bottom, the voltage of the model, gating variables *a* (black) and *b* (red), SIK current *I_SIK_*. (B) Simulation of the two current steps *I_inj_* = 188*pA* with a resting time Δtime = 200*ms*. From top to bottom, injection current, gating variables *a* (black) and *b* (red), the voltage of the model. (C) The peak SIK current and delay of the first spike recover with a long resting time Δtime. The recover ratio of the peak SIK current (red) is calculated by comparing the maximum *I_SIK_* during the second voltage step to that during the first voltage step. The delay of the 1st spike is calculated from the current steps (black).

### Onset bursting is reproduced in an *α*7 cell model

We then studied the onset bursting behavior, which is observed in most of the *α*7 cells. Cells with an OB feature show a characteristic high onset firing rate and a fast dropping of the firing rate during the first 40*ms* (Figure 7A). In the phase diagram of the cell, several lines are above the spiking cycle (red arrow in Figure 7B), which indicates the existence of a transient current that is less dependent on the voltage. Further, the onset firing rate is about 270*Hz* across sweeps with different injection currents. The firing rate robustly drops below 50*Hz* within 40*ms* (Figure 7C).

**Figure 7.**
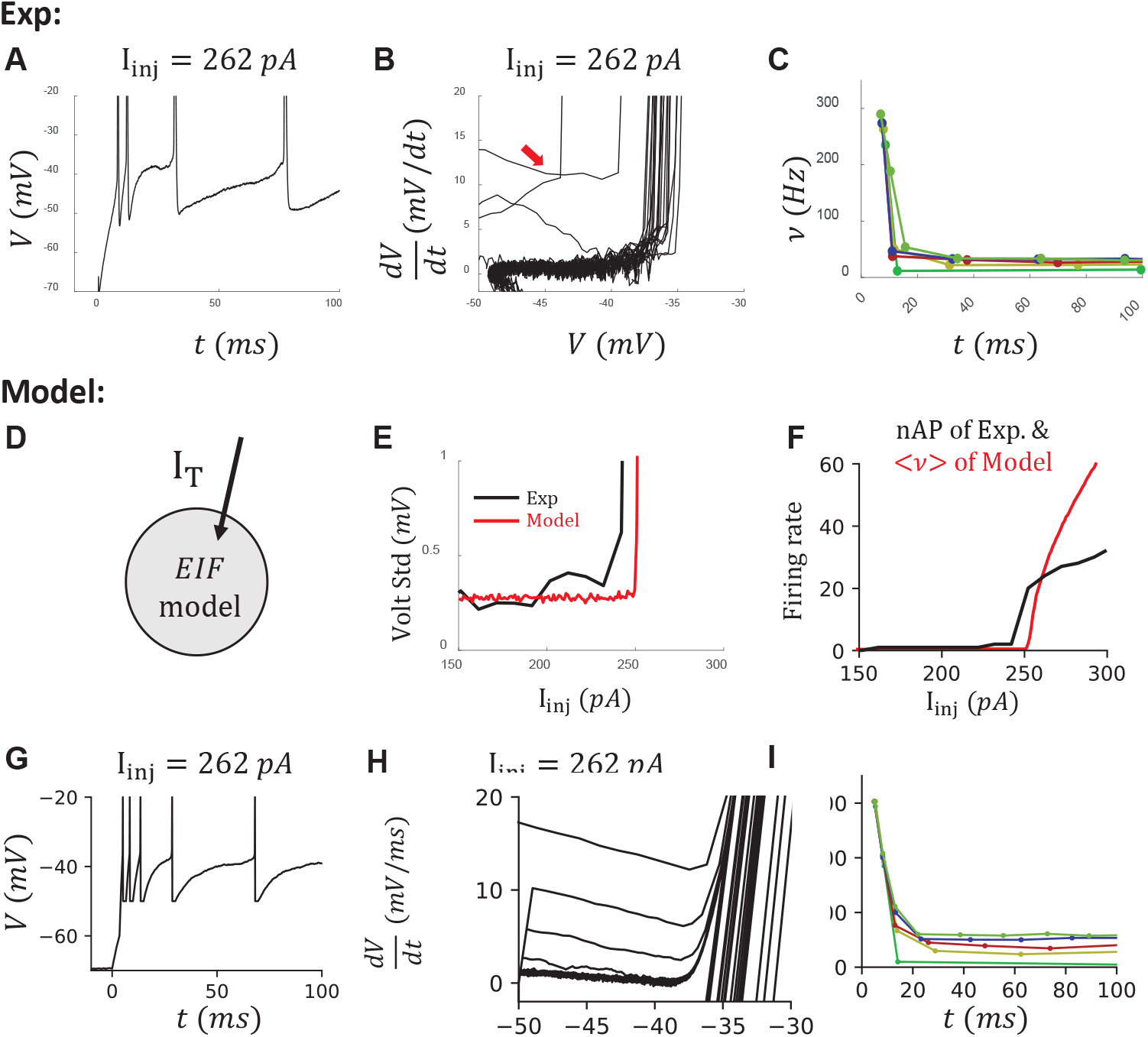
Modeling the onset bursting of an *α*7 cell with a T-type calcium channel. (A) The voltage trace at *I_inj_* = 262*pA*. (B) Phase diagram of the cell. The red arrow indicates the trajectory from the first AP within the onset bursting period. (C) Instantaneous firing rate shows a sudden drop in the first 40*ms* of the injection period. Colors suggest sweeps with different injection currents *I_inj_* = 252, 262, 272, 282, 292*pA*. (D) The sketch of the *α*7 cell model. (E) The voltage standard deviations of the data and the model. (F) The firing rates from the data and the model. (G to I) Modeling results organized as (A to C).

**Figure 8.**
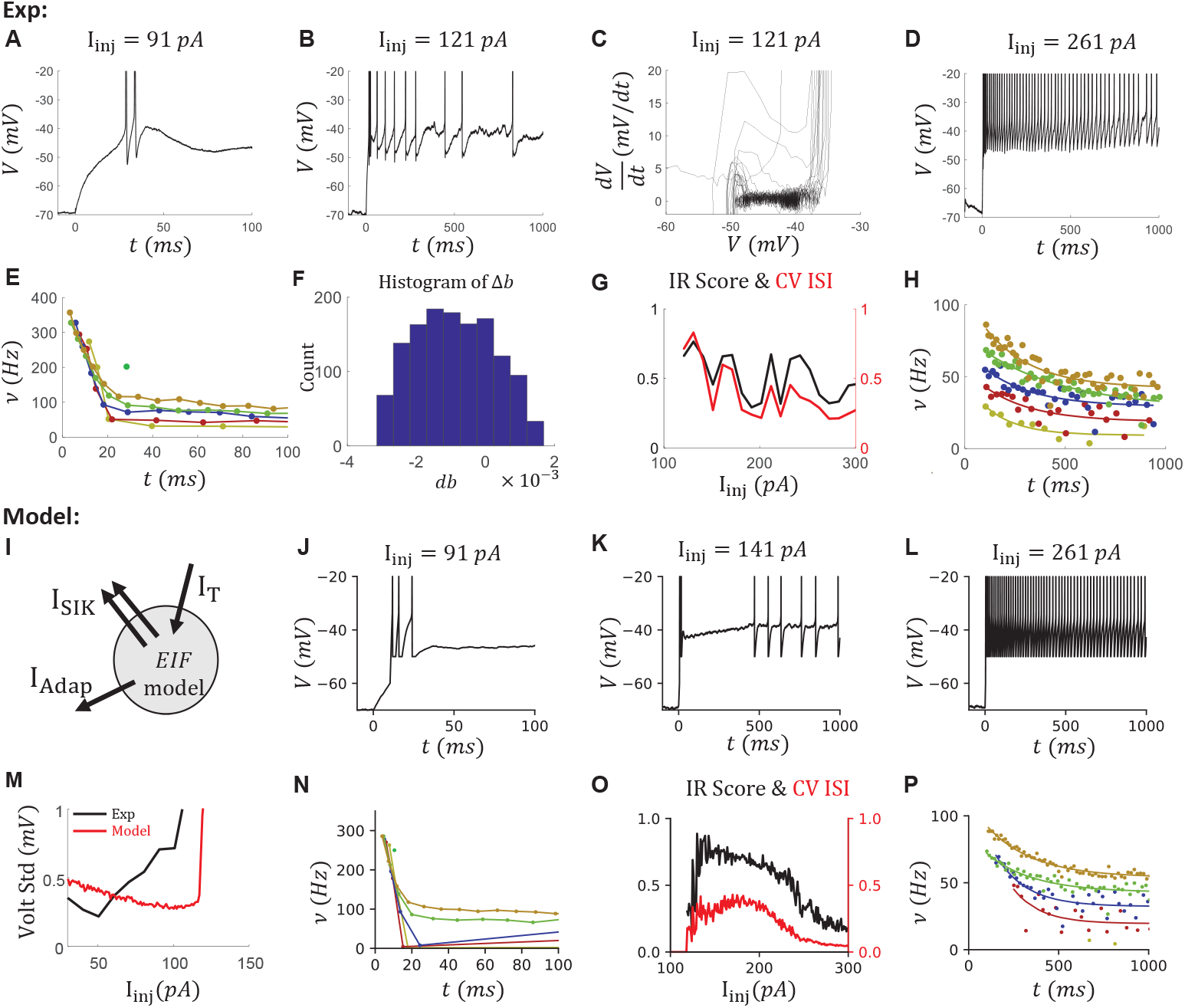
Reproducing the onset bursting, irregular firing patterns, and adaption simultaneously in a VIP cell model. (A to D) Voltage traces at *I_inj_* = 91, 121, 261*pA*, which are examples to show OB, IR, and Adap, respectively. (E) The instantaneous firing rates *ν* from 0 to 100*ms* that show the onset bursting. Colors represent sweeps with different injection currents *I_inj_* = 91, 131, 171, 212, 251, 291*pA*. (F) The histogram of Δ*b* assuming *b_init_* = 0.25 just before the AP. The average change is < Δ*b* >= −0.0009. (G) The ISI ratio (black) and CV ISI (red) curves that show the irregularity. (H) The instantaneous firing rates *ν* between 100*ms* to 1*s* that show the adaptation. Color coding is the same as (D). (I) Sketch of the VIP cell model. (J to P) Corresponding Modeling results. (M) shows the standard deviation of the voltage of the data and the model.

As previously suggested (Schuman et al., 2019) and demonstrated in (Schuman et al., 2021), OB in *α*7 cells is depended on the expression of a T-type *Ca*^2+^ channel. This was also demonstrated in bursting L2/3 VIP cells (Prönneke et al., 2020). We thus include a T-type *Ca*^2+^ current in our EIF model for an *α*7 cell model (Figure 7D) based on the reported dynamics of T-type *Ca*^2+^ channels from a thalamocortical relay neuron model (Smith et al., 2000). After tuning the T-type *Ca*^2+^ channel conductance *g_T_* to the OB in the data, along with noise level *σ* (Figure 7E) and other passive parameters, we can reproduce the high onset firing rate and the fast dropping of the firing rate during the first 40*ms* in the *α*7 cell model (Figure 7F to I).

### IR, OB and Adap are simultaneously reproduced in a VIP cell model

After reproducing the IR, Acc, and OB features individually, we turn to the complex dynamics observed in VIP cells. Most VIP cells show more than one considered features (Figure 1B4). For example, onset bursting (Figure 8A, E), irregularity (Figure 8B, G), and adaptation (Figure 8D, H) are observed in the same selected VIP cell. All these features can be simultaneously reproduced with an EIF model with a T-type *Ca*^2+^ channel, a SIK channel, and a spike-triggered small conductance *K*^+^ channel (Figure 8I).

As described in the modeling of an *α*7 cell (Figure 7), the OB has the characteristic high onset firing rate and a fast dropping in the firing rate during the first 40*ms*. This was reproduced by tuning the conductance *g_T_* of the T-type *Ca*^2+^ channel, as in the *α*7 model.

In addition, the IR score is highest in the sweeps around rheobase, and it decays slowly with increasing injection current (Figure 8G). The IR score remains higher than 0.4, which is the threshold to detect IR, even in a 1*s* sweep with 60 APs with injection current *I_inj_* = 300*pA*. Above, we reproduced IR spiking by including a strong SIK current in the Canopy cell model. However, the IR score drops fast in the Canopy cell model when increasing the injection current (Figure 3K).

Further, strong adaptation is also observed in the VIP cell in the last 900*ms* (Figure 8D), where the firing rate drops exponentially to a saturation value (Figure 8H). To quantify that, we define the adaptation index (AI) as the fraction of the firing rate that drops during this 900*ms* period. For this VIP cell, AI is about 0.5 in all the sweeps (Figure 9E). To reproduce the adaptation, we include a spike-triggered small conductance *K*^+^ channel in our model (Figure 8I) that generates after-hyperpolarization (AHP) current reproducing current-independent adaptation, as in (Liu and Wang, 2001).

**Figure 9.**
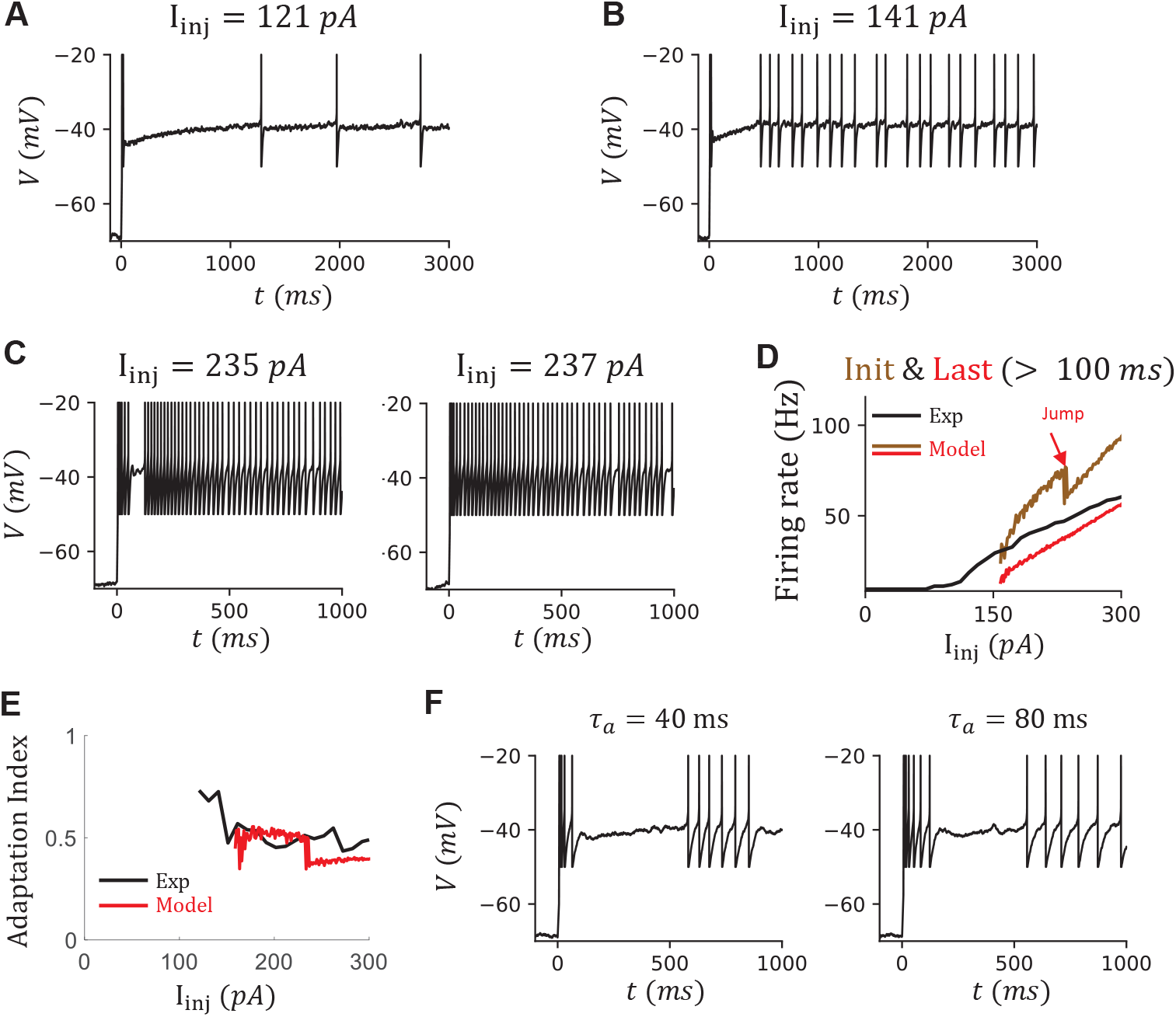
More results of the VIP cell model. (A, B) The voltage traces of the model in 3*s* at *I_inj_* = 121, 141. (C) The voltage traces of the model at *I_inj_* = 235, 237*pA*. The jump in the curve of the initial firing rate is due to the diminishing of the silent period after the onset bursting. (D) The comparison of the firing rates *ν* of the data and the model. The *ν* of the model is measured as the number of APs during 1s current injection. The initial and last (steady-state) firing rates *ν* of the model are measured from 100*ms* to 1500*ms*. (E) The adaptation index (AI) from the data (black) and the model (red), measured based on the fitting of 1/ISI from 100*ms* to 1*s*. (F) Voltage traces when varying the time constant of fast activation variable *τ_a_* = 40, 80*ms* with an injection current *I_inj_* = 141*pA*. The firing pattern around 100*ms* is more comparable between the data and the model. However, *τ_a_* is not biologically plausible anymore (Original *τ_a_* = 4*ms*).

After a similar tuning of the VIP cell model, based on the phase diagram (Figure 8C), the noise level (Figure 8M), the jump of the Δ*b* (Figure 8F), we can reproduce simultaneously the OB, which is reflected in the high onset and the fast-dropping of the firing rate during the first 40*ms* (Figure 8J, N), IR, which is reflected by the high IR score and high CV of the ISIs (Figure 8K, O), and Adap, which shows exponentially decay during the latter 900*ms* (Figure 8L, P).

However, there are limitations to this VIP cell model. First, though the irregularity during the steady-state is similar between the data and the model, the onset of irregular firing in the model is usually later than that in the data. Especially when the injection current is around the rheobase, the irregular firing may be observed after 1*s* in the model (Figure 9A). In contrast, in the data, it may be observed between 500*ms* to 1*s* (compare Figure 8B). Second, the cell in the model becomes quiescence after the OB period (around 100*ms*, Figure 9B, C), which results from the strong adaptation current triggered by several APs during the onset bursting and the fast activation of the SIK current. But this is not the case in the data (Figure 8B). In the model, this quiescence period diminishes when the injection current is high enough inj= 235*pA* (Figure 9C). Since we are measuring the adaptation starting at 100*ms*, the diminishing of the quiescence period is reflected in the onset firing rate after 100*ms* (Figure 9D) and is further reflected in the jump on the AI curve (Figure 9E).

To get an insight into what impacts firing after the OB period, we change the timescale of the fast activation variable of the SIK channel, which delays the onset of outward SIK current (Figure 9F). As expected, the firing after the OB continues. But to achieve a comparable length with the data, the timescale needs to be around 80*ms*, which is unlikely for a SIK channel. We discuss this discrepancy further in the discussion section.

### Mechanism of irregular firing in the VIP cell model

To investigate the origin of IR in the VIP cell model, we do a similar fast-slow analysis as we did for the canopy cell model. We first exclude the noise term in the model by setting *σ* = 0 and observe cluster spiking in the simulation (Figure 10A). In contrast to the canopy cell model, we have two slow variables: slow inactivation variable *b* and the dimensionless *Ca*^2+^ concentration *C_ca_*. To do the fast-slow analysis, we assume both are constants. Similarly, the system has one SS and one US when either outward current prevails the injection current (Figure 10B). By assuming *C_ca_* is a constant, we can define the *b_min_*(*C_ca_*) as the minimal *b* where a global SS exists (Figure 10C, red dashed lines). In the full system, the line *b_min_*(*C_ca_*) represents the line where the SNIC bifurcation happens (Figure 10D, red dashed lines). Only in the region above the bifurcation line, a global SS exists. In the simulation of the full model (Figure 10A, D), the system oscillates between a resting state and a spiking state, which is represented as cluster spiking. The dynamics change to irregular firing when adding noise to the system by setting *σ* > 0.

**Figure 10.**
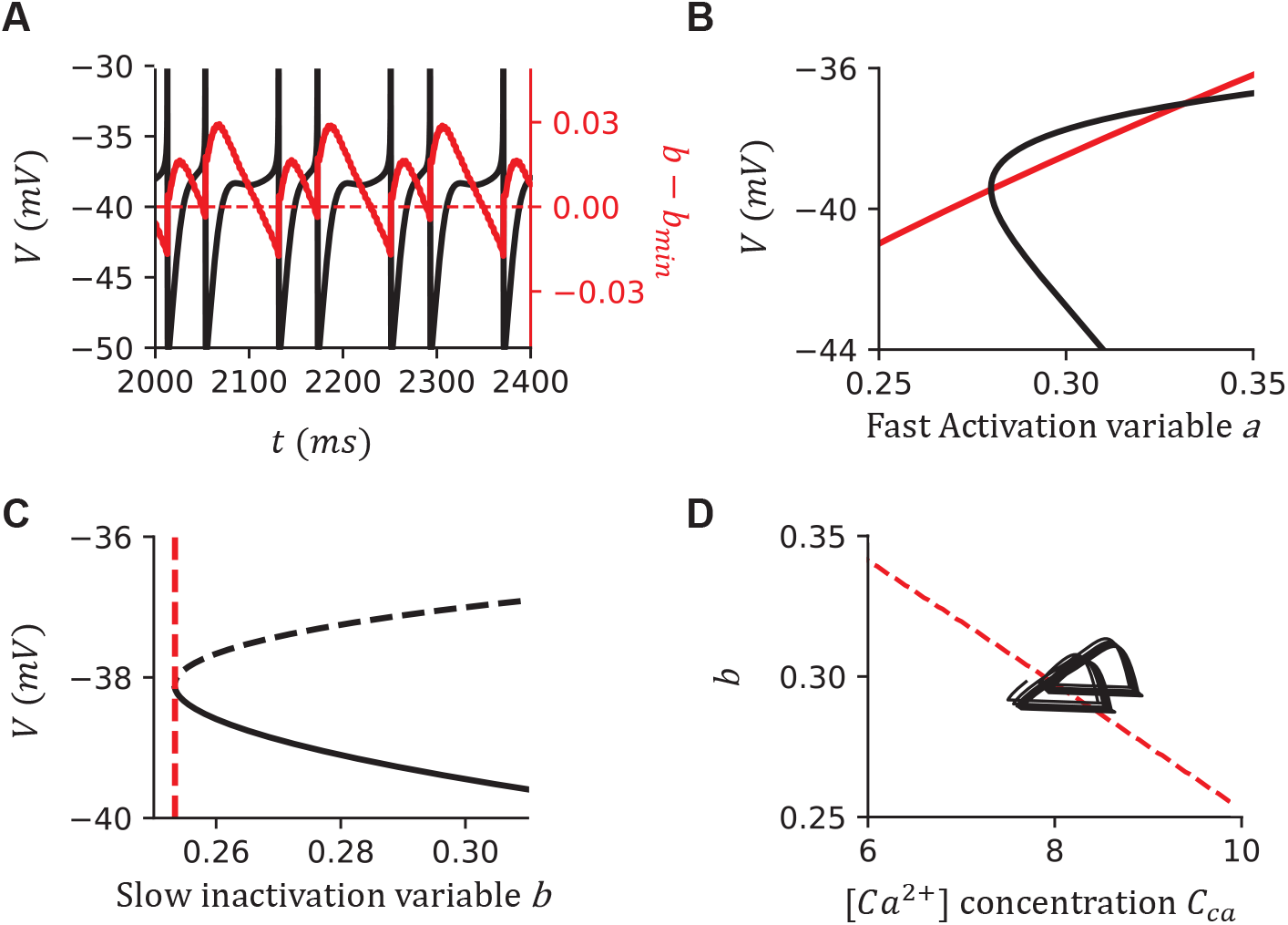
The fast-slow analysis of the VIP cell model shows that the irregularity results from the square-wave bursting. (A) The sample simulation when *I_inj_* = 170*pA*. The black line indicates the voltage trace, and the red line indicates the difference between *b* and *b_min_*. The red dashed line is at *b − b_min_* = 0, suggesting the boundary between the resting state and the firing state. (B) Nullclines of the fast manifold when *I_inj_* = 170*pA*, *C_ca_* = 10, *b* = 0.3. The intersections indicate the fixpoints of the fast manifold. (C) The bifurcation diagram when *I_inj_* = 170*pA*, *C_ca_* = 10. The black solid and dashed lines indicate the stable branch and the unstable branch, respectively. The red dashed line indicates the minimum of *b* that the fast manifold has a global SS. (D) The bifurcation diagram of the whole system and the simulation when *I_inj_* = 170*pA*. The red dashed line *b_min_*(*C_ca_*) indicates the SNIC bifurcation line. Only the region above which has a global SS. The black line indicates the simulation of the VIP cell model without noise (*σ* = 0).

**Figure 11.**
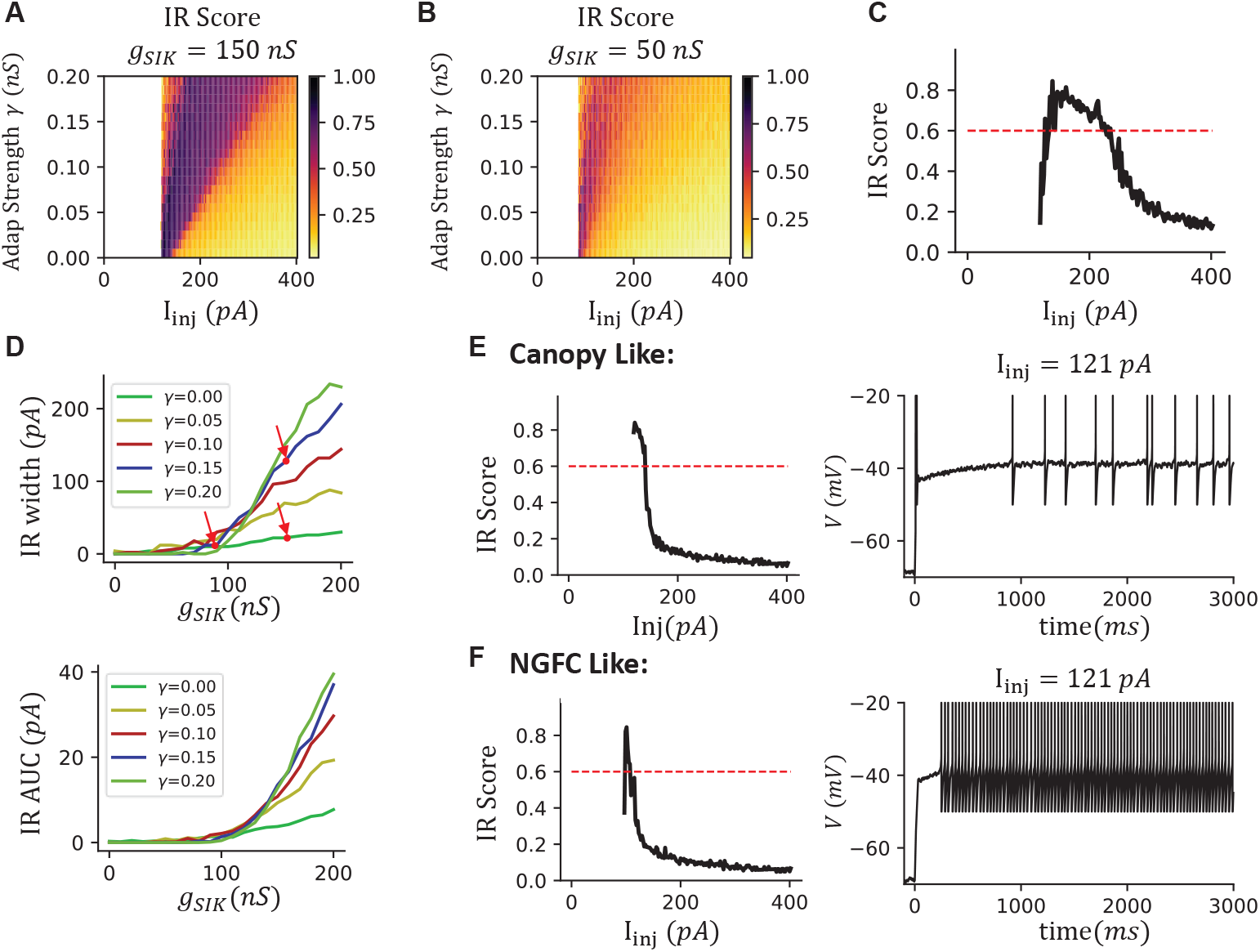
The irregular firing of VIP cells is more easily observed in experiments because of a large irregular width introduced by the interaction between *I_SIK_* and *I_adap_*. (A) The ISI ratio (color) over different injection currents and adaptation strength *γ* when *g_SIK_* = 150*nS*. (B) as (A) but *g_SIK_* = 50*nS*. (C) The sample ISI ratio when *g_SIK_* = 150*nS, γ* = 0.1. The range of injection current when CV ISI is above the threshold 0.6 is defined as the irregular width (IR width). (D) The irregularity visibility is high only when the *g_SIK_* and *γ* are both large. Top: The IR width curves over *g_SIK_*. Different lines suggest results with different adaptation strength *γ*. The dots and arrows indicate the parameters of the VIP cell model (top-right), the Canopy-like case (bottom-right, shown in E) and the NGFC-like case (left, shown in F). Bottom: the IR area-under-curves, defined as the area under the curve of ISI ratio and above the ISI ratio threshold 0.6. (E) The model results when set *γ* = 0, *g_SIK_* = 150*nS* look like a canopy cell. (Left) The IR score. The drop of the curve is sharp when increasing the injection current, and the IR width is small. (Right) Voltage trace at *I_inj_* = 121*pA*. (F) The model results when setting *γ* = 0, *g_SIK_* = 90*nS* looks like a NGFC. The IR width is further reduced in comparison to (A). In this case, we remove the T-type *Ca*^2+^ current for better visualization.

**Figure 12.**
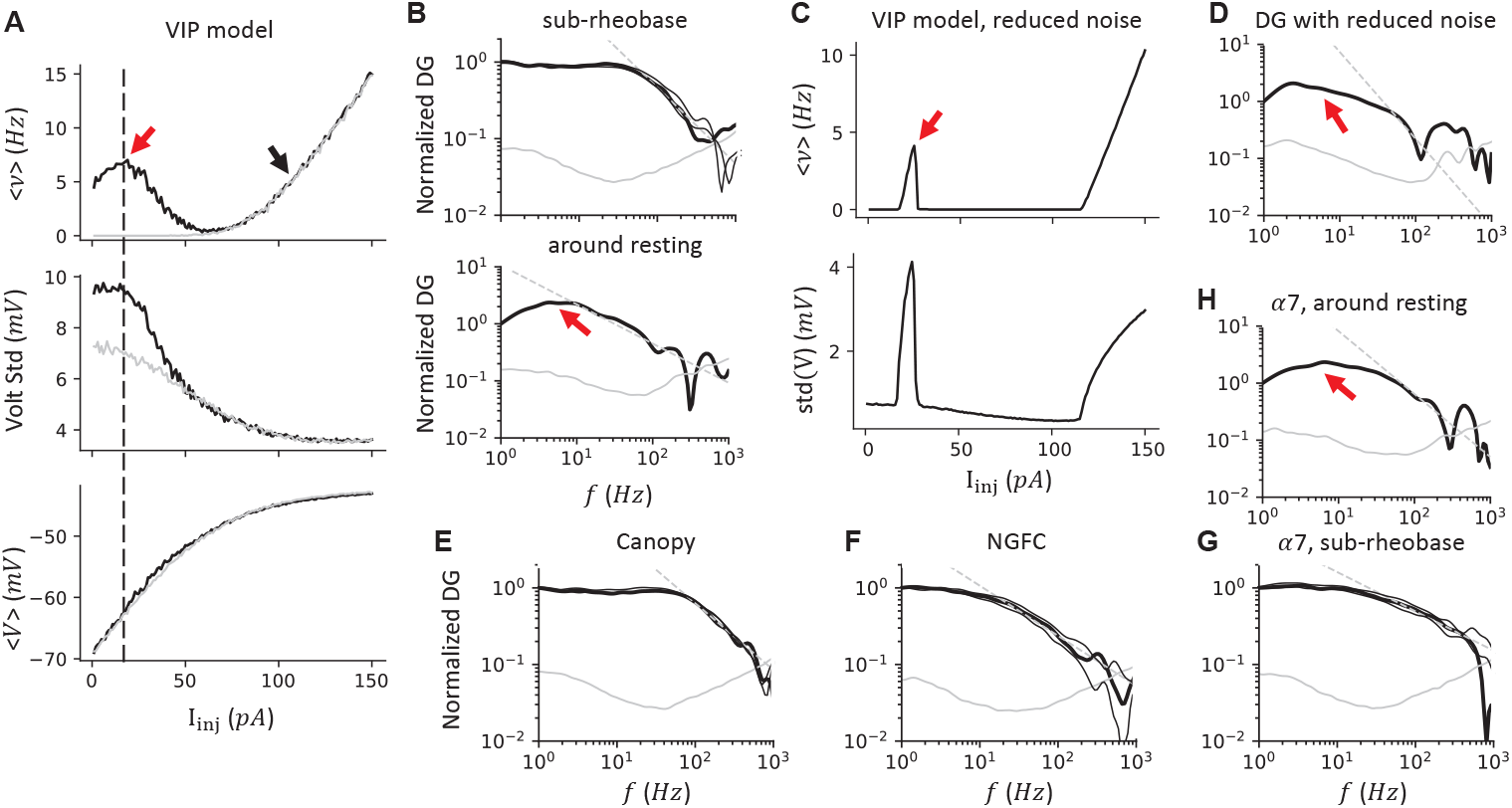
The VIP model resonates with Theta/Alpha band input only when the average input is around the resting. (A) Simulation results of the VIP model when varying the average injection current. In this panel, *σ* = 50*pA*. Gray lines, model results while excluding the T-type *Ca*^2+^ channels. Top, the average firing rate *ν* shows two non-zero branches. Mid, the voltage’s standard deviation decreases when the injection current increases. Bottom, the average voltage of the model. (B) The normalized dynamic gain when the injection current is at the sub-rheobase region (*I_inj_* = 20*pA*, black arrow in A) and around the resting (*I_inj_* = 105*pA*, red arrow in A). The DG shows an unexpected high-gain range around the Theta/Alpha band when the model is around the resting regime. The high-frequency profile was described as *f ^−α^* with a cut-off frequency *f*_0_ and *α*, shown in a dashed gray line. The two thin-black lines represent the simulation with average firing rates < *ν* >= 3, 7 Hz, respectively. The solid gray line indicates the 95% confidence level generated by bootstrapping the shuffled ISIs. Top, *f*_0_ = 58.9*Hz*, *α* = 0.99; bottom, *f*_0_ = 67.6*Hz*, *α* = 0.69. (C) Simulation results of VIP model with reduced noise *σ* = 5*pA*. Top, the average firing rate; bottom, the standard deviation of the voltage. (D) The DG around the resting still shows a high gain range around the alpha band (*I_inj_* = 25*pA*, red arrow in C) when the input noise is drastically reduced. *f*_0_ = 41.7*Hz*, *α* = 1.52 (E to G) The normalized DG for the Canopy cell model (*I_inj_* = 282*pA*,*σ* = 90*pA*, *f*_0_ = 77.6*Hz*, *α* = 0.62), NGFC model (*I_inj_* = 170*pA*,*σ* = 80*pA*, *f*_0_ = 14.8*Hz*, *α* = 0.65) and *α*7 cell model (*I_inj_* = 200*pA*,*σ* = 65*pA*, *f*_0_ = 38.9*Hz*, *α* = 0.51) when the average input current is at sub-rheobase. (H) The *α*7 cell model also has a non-zero firing branch around the resting that shows high gain around the Theta/Alpha band frequency(*I_inj_* = 25*pA*,*σ* = 65*pA*, *f*_0_ = 83.2*Hz*, *α* = 1.10).

### Mechanism behind the traditional alias “irregular spiking cell” for VIP cells

Traditionally, the term irregular spiking has been used mainly for VIP cells but rarely for other INs (Tremblay et al., 2016). But, in our study, the irregularity is not uniquely found in VIP cells but distributed in multiple subtypes of L1 interneurons. Especially, the IR in the VIP cells is not due to a higher noise level. In fact, the noise levels around the rheobase are comparable across all the subtypes (Figure 3G, 5F, 7E, 8M). Is there anything unique for the IR VIP cells but not other INs with an IR feature?

To answer this question, we systematically vary the parameters of our VIP cell model. We measure the IR score while varying adaptation strength *γ* and the SIK conductance *g_SIK_* at different injection currents (Figure 11A, B). The IR score is higher when the SIK conductance is bigger. Interestingly, the high IR score is more spread out with a stronger adaptation. To quantify this effect, we define IR width as the range of injection current where the IR score is bigger than the threshold 0.6 (Figure 11C). As a result, we observed that the IR width only starts increasing when *g_SIK_* reaches around 100*nS* (Figure 11D, top). However, the IR width remains small if an adaptation current is not included *γ* = 0. The IR width only becomes large when the adaptation strength is also increased. The results are similar if we use the IR area-under-curve (AUC) instead of IR width (Figure 11D, bottom). These explain the behavioral differences observed in the Canopy, NGFC, and VIP models.

In our VIP cell model, the IR width is around 120*pA* (right top arrow in Figure 11D, *g_SIK_* = 150*nS*, *γ* = 0.1). When the adaptation current is removed by setting *γ* = 0, the IR width decreases drastically to around 20*pA* (right bottom arrow in Figure 11D), where the dynamics mimic the IR in Canopy cells (Figure 11E). If in addition we decrease SIK conductance to *g_SIK_* = 90*nS*, the IR width decreases to about 10*pA* (left arrow in Figure 11D), where the dynamics mimic the Acc in NGFC (Figure 11F). These results suggest that IR is most easily observed only when the *g_SIK_* is big and the adaptation is strong, which is the case for VIP cells but not other INs.

To verify this claim in our dataset, we measured the IR width by multiplying the number of sweeps with an IR score bigger than 0.4 by the corresponding current step size of IR cells in different IN subtypes. The average IR width of VIP cells is the highest, and it is much higher than that of canopy cells (Table 2). Furthermore, the input resistance is the highest of the VIP cells compared to other L1 INs (Table 2), including *α*7 cells. Even though the IR width of VIP cells and *α*7 cells are comparable, the corresponding voltage change of VIP cells is much higher than that of *α*7 cells, which explains why the IR feature is most salient in VIP cells.

### VIP cell model resonant with Theta/Alpha input through the activation of T-type *Ca*^2+^ channel

Due to the intrinsic properties, neurons may respond differently to an input of varied frequencies though the mean is the same. Theoretical and experimental studies suggested that these responses, which are referred to as the dynamic gain (DG), are important in understanding the response from a neuron ensemble (Fourcaud-Trocmé et al., 2003; Geisler et al., 2005; Köndgen et al., 2008).

Recently, (Borda Bossana et al., 2020) reported rat L1 INs are homogeneous and narrow-tuned through the DG analysis by using a colored noisy input generated by an Ornstein-Uhlenbeck (OU) process, developed by (Higgs and Spain, 2009) (See also (Ilin et al., 2013)). Briefly, in this analysis, a subthreshold colored noisy current signal, generated from an OU process with a time constant *τ_σ_* = 5*ms*, is given as input. Since the average input *I_inj_* is subthreshold, the responding APs are results of noise, but not a tonic, component of the input. The average current *I_inj_* and noise level *σ* are tuned such that the average firing rate is about < *ν* >= 5*Hz* and the standard deviation of voltage is about 4*mV*. These procedures are required to have similar membrane potential fluctuations that recorded *in vivo* (Ilin et al., 2013). Next, long recordings are made (at least 1000 seconds long and collecting 5000 APs), and further analyzed in the Fourier space. The ratio between the response and the input at a specific frequency is defined as the dynamic gain at that frequency (*DG*(*f*). referred to as the dynamic transfer function in (Borda Bossana et al., 2020)).

Here, we test the response of our VIP cell model, along with other L1 IN models, to the different rhythm inputs through the dynamic gain analysis. We find that with a designed synaptic input fluctuation (tuned at the black arrow in Figure 12A), the VIP cell shows two non-zero firing rate branches (Figure 12A). The sub-rheobase branch is expected where the APs are triggered by colored noise. However, when we decrease the average injection current while keeping the noise *σ* the same, the second branch around the resting voltage emerged, with an increased potential fluctuation and a peak around −60*mV*. This second branch disappears if the T-type *Ca*^2+^ channel is removed from the model (*g_T_* = 0, Figure 12A, gray lines). Within the sub-rheobase regime (Figure 12B, top), the DG shows a similar “narrow bandwidth” as reported in (Borda Bossana et al., 2020). The results are robust when the average firing rate varies (3 and 7*Hz*, Figure 12B, top, two thin lines). Interestingly, around the resting regime (Figure 12B, bottom), the model resonates to the Theta/Alpha (4 − 15*Hz*) band input. The observed resonance has characteristics that agree with the dynamic of the T-type *Ca*^2+^ channel. This resonance happens at the average voltage around −60*mV*, where the T-type *Ca*^2+^ channel activates (Figure 12A, bottom). Also, the Alpha/Theta rhythm matches the activating time constant of the T-type *Ca*^2+^ channel 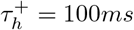.

To make the voltage fluctuation comparable, we reduced the input noise *σ*, while the non-zero firing branch around the resting remains (Figure 12C). Similarly, we observed high gain to the Theta/Alpha band input (Figure 12D).

We further tested the DG in other IN models (Figure 12E to H). All model shows a similar “narrow bandwidth” feature at the sub-rheobase regime, suggesting the DGs of L1 IN models are homogeneous, as reported in (Borda Bossana et al., 2020). Only the *α*7 model, which includes the T-type *Ca*^2+^ channel, shows another non-zero firing regime around the resting, where the model resonates to the Theta/Alpha band input (Figure 12H).

## Discussion

The neocortex is characterized by a remarkable diversity of GABAergic interneuron subtypes (Markram et al., 2004). Their functions have been proposed to serve diverse functions, exemplified by the well-established disinhibitory motif composed of three inhibitory subtypes (Wang et al., 2004; Tremblay et al., 2016). In this study, we systematically analyze the distribution of electrophysiological features in the L1 IN subtypes and further build models for each subtype by including different ion channels. Canopy cells and NGFCs mostly show irregular firing or acceleration in the firing rate, suggesting SIK exists in these two cell types. Most of the *α*7 cells are classified as having an onset bursting feature, which implies the presence of a T-type *Ca*^2+^ channel. The VIP cells are very heterogeneous, but many show adaptation and irregular firing. To reproduce that, we build a model that includes a SIK channel, a T-type *Ca*^2+^ channel, and a spike-triggered *Ca*^2+^-dependent *K*^+^ channel. We apply slow-fast analyses to show that the irregularity comes from the square-wave bursting, which requires a strong SIK current. Further, we showed that the adaptation current observed within the VIP cells significantly expands the region where the irregular firing can be observed, and this is the main reason why VIP cells are often considered “irregular spiking cells” but not other INs (Tremblay et al., 2016).

Our work provides insights into outstanding questions concerning interneuron subtype biology. First, by revealing the mechanism of IR firing of VIP cells, applying an antagonist to either block the SIK channel or the adaptation current should eliminate the irregular firing. Applying gene-editing methodology in cultured cells should show similar results as well. Second, our VIP model suggests that the different electrophysiology subtypes from VIP cells may come from continually varying the strength of different ion currents, thus contributing to the classification of VIP subtypes. For instance, the cells classified as IS, fAD, and bNA in (He et al., 2016) may be reproduced by just including a strong SIK current, a strong T-type *Ca*^2+^ channel with or without a strong adaptation current, and a combination of T-type *Ca*^2+^ current with a SIK current, respectively. Third, we revealed an unexpected resonance of Alpha/Theta band input in the VIP cell model and *α*7 cell model around the resting voltage, which can be measured experimentally. Previously in (Higgs and Spain, 2009), the bursts generated from the fast afterhyperpolarization provide a high gain at a similar rhythm band (7 − 16*Hz*). Similarly, our model predicts that the Alpha/Theta resonance is associated with the onset bursting, and both features arise from the dynamics of the T-type *Ca*^2+^ channel.

Detailed reconstructed models of rat L1 INs, along with every neuron subtype, were studied in the Blue Brain Project (BBP, (Markram et al., 2015)) through a systematically optimizing algorithm. Within their study, most L1 INs are classified as continuous non-accommodating cells (cNAC), while only a few are stuttering (cSTUT) or irregular firing (cIR). However, only a limited number of sweeps (1.5×, 2×, 2.5× rheobase) are used in classification and further optimizing their models. In our dataset, canopy cells and NGFCs account for about 70% of the total L1 INs population, and most are classified as having an IR or an ACC feature. This does not contradict the BBP observation since we consider the finer recordings around the rheobase. If we only consider the recordings away from the rheobase, the NFGCs and Canopy cells would be classified as cNACs as BBP did.

Further, the BBP models and our model differ in reproducing irregular firing. The BBP models introduce the high channel noise from a stochastic potassium channel to reproduce the irregularity ((Markram et al., 2015), Supp. Materials, Optimization of neuron models). However, we showed that this is unnecessary if a SIK channel is included. In fact, our data shows that the noise level is comparable between irregular firing cells and others (Figure 3G, 5F, 7E, 8M), which suggests that high channel noise is not the reason for the irregular firing.

We build our models based on an EIF model with a few additional ion channels to reproduce the major observed dynamics. Thus, the models are easy to analyze, and the computational cost is minimized when implemented in a large-scale neuronal circuit, contrasting to the detailed reconstructed multicompartmental models (see (Markram et al., 2015) for examples). However, doing so limits the ability to reproduce electrophysiological features from various aspects. Unlike Hodgkin-Huxley type models, our model doesn’t include the dynamics during the APs. Thus we cannot reproduce the features of APs from different subtypes (see Table 2) in our model. Another issue is that we mostly reproduce the firing behavior around the rheobase (< 20*Hz*), but not with high firing rates. This is because our dataset includes recordings mostly around the rheobase but not with high firing frequency (most recordings have < 30 APs), and our EIF model, unlike the Hodgkin-Huxley type model that stops firing when the outward currents are overwhelmed by the injection current, can have arbitrary high firing rates. In addition, the f-I slope in our model is higher than that measured from the data. From our experience, only the capacity significantly impacts the f-I slope in the model, which is constrained (Figure 2, Table 2). We speculate that this mismatch can be alleviated by including other outward currents, like *K*^+^ current from KCNQ channels (Goff and Goldberg, 2019).

Late-spiking is a signature of neurogliaform cells, but the underlying mechanism is unknown. In delayed-firing hippocampal CA1 pyramidal neurons (Storm, 1988), blocking Kv1, a SIK current, with low concentrations of 4-aminopyridine (4-AP), eliminates the delayed-firing feature. Further, in delayed fast-spiking cells, a dendrotoxin sensitive Kv1 current underlies the delayed firing ((Goldberg et al., 2008), see also (Bos et al., 2018)) and clustered spiking with subthreshold oscillations (Sciamanna and Wilson, 2011). Based on these experiments, we include a SIK channel in our NGFC model and reproduce the late-spiking and firing-rate acceleration. However, a recent study on NGFCs suggested that even though Kv1 currents exist in the mouse NGFCs and contribute to the delayed-firing ((Chittajallu et al., 2020) Figure 4I, applying low 4-AP decreases about 65% of the AP latency), their functional role may be lesser than that of other *K*^+^ currents, especially from the Kv4 family. Furthermore, preliminary results from our lab suggest that dendrotoxin does not block delayed firing in NGFCs. Interestingly, the recovery time from inactivation is about 1*s* in (Chittajallu et al., 2020) and about 12*s* in (Bos et al., 2018), which is slower than that in our model (about 500 ms, Figure 6C). Nevertheless, this discrepancy is not likely explained by the dynamics of Kv4 channels, which are faster than that of Kv1 channels (Coetzee et al., 1999). Further pharmacological experiments are necessary to help discover the slowly inactivating *K*^+^ current of NGFCs.

A significant difference between NGFCs and Canopy cells is that NGFCs are late-spiking whereas Canopy cells are not. However, if Canopy cell models only include a SIK channel, they are also late-spiking (Figure 3M). Also, VIP cells often show decreased firing rates after the OB period, usually around 100*ms*, but are not quiescent (Figure 8B). This behavior was hard to reproduce in the model, due to the large adaptation current triggered by several APs in the OB period. These discrepancies in the Canopy model and the VIP model suggest the existence of transient inward currents (or inactivation of outward currents) that have slower dynamics than the T-type *Ca*^2+^ current. In VIP cells (Goff and Goldberg, 2019), the Nav1.1 channel may contribute to the length of the onset firing period via progressive inactivation during repetitive firing. Further studies are required to validate the role of Nav1.1 channels in models with an IR feature.

Recently, it has become clear that L1 is the key to understanding how long range cortical-cortical interactions alter local circuit dynamics (Schuman et al., 2021; Ledderose et al., 2021). For example, recent work in rodents showed that hippocampal-dependent associative learning is controlled by perirhinal input to L1 (Doron et al., 2020; Shin et al., 2021). More importantly, this associative learning could be diverted by dendritic inhibition. Another recent study had shown that L1 NDNF+ INs (NGFCs and Canopy cells in our study) can shift the input gain from dendrite to soma in pyramidal cells across all layers in the cortex (Malina et al., 2021), suggesting L1 NDNF+ INs may be well-suited to control associative learning. Further, another anatomical study showed that ventromedial and mediodorsal thalamus target NDNF+ cells and VIP+ cells in mouse prefrontal cortex, respectively (Anastasiades et al., 2021), through which these higher-order thalamic nuclei differentially modulate the neuronal activity in the prefrontal cortex. To further investigate the hypotheses listed here and beyond, a well-developed model that incorporates the distinct properties of each L1 interneuron subtype will help in elucidating the mechanisms by which the L1 circuit enables the integration of top-down and bottom-up information streams, and will assist in the generation of predictions to guide future experiments. Our work on the individual L1 IN subtypes here is an essential step towards this goal.

## Acknowledgments

This work was supported by National Institutes of Health grants R01NS110079 to B.R. and X.W., P01NS074972 to B.R., and R01MH062349 to X.W. The authors thank Alvar Prönneke, Sean Froudist-Walsh, Kevin Berlemont and members of the Wang and Rudy labs for discussion.

## Extended Data

Table 1-1 The analysis on individual L1 INs.

